# Replicated radiations reveal temporally structured macroevolutionary dynamics

**DOI:** 10.1101/2025.05.13.653847

**Authors:** Callum V. Bucklow, Emanuell Duarte Ribeiro, Fabrizia Ronco, Nathan Vranken, Michael K. Oliver, Walter Salzburger, Melanie Stiassny, Mexford Mulumpwa, Roger Benson, Berta Verd

## Abstract

Macroevolutionary theory predicts that adaptive radiations are characterised by early bursts of trait evolution followed by rate slowdowns as ecological niches are filled. However, these dynamics are rarely detected within individual clades. Here, we test whether this discrepancy arises from limited temporal sampling by comparing replicated adaptive radiations of African cichlids spanning distinct evolutionary timescales. We show that the net rates of vertebral count evolution are inversely correlated with radiation age, consistent with early burst dynamics. However, we do not detect this pattern within the individual systems. We also find evidence of changes to the underlying processes that generate disparity across the timescale of adaptive radiation, signalled by increasing importance of phylogenetic signal with radiation age. This suggests that early phenotypic expansion is both rapid and phylogenetically unstructured, but that stabilising dynamics emerge on longer timescales, amid declining rates. Despite rapid phenotypic diversification, lacustrine lineages occupy only a subset of the broader morphospace of riverine taxa, suggesting that diversification over short timescales is constrained and may largely resample the variation already present in older lineages. Our results provide strong evidence for shifting evolutionary dynamics through time in adaptive radiations, and that the apparent rarity of early-burst signals in comparative analyses may partly reflect limited statistical power due to narrow temporal sampling.

## INTRODUCTION

Adaptive radiation is the rapid diversification of lineages following expansion of ecological opportunity, during which species proliferate, and phenotypic disparity accumulates (Simpson, 1953; Schluter, 2000; Stroud and Losos, 2016). Early phases of radiation may have high rates of trait evolution, rapid variance accumulation, and shifts toward distinct adaptive optima (Simpson, 1953; Harmon et al., 2010; Etienne and Haegeman, 2012; Slater and Friscia, 2019). Over time, as ecological niches are filled, evolutionary dynamics are expected to transition toward slower rates and stabilising processes consistent with movement around established adaptive peaks (Butler and King, 2004; Mahler et al., 2010). Despite these strong theoretical expectations, key macroevolutionary signatures such as early bursts of trait evolution and subsequent rate slowdowns are rarely detected within individual clades (Harmon et al., 2010; Slater and Pennell, 2014), suggesting either that such dynamics are uncommon or that they are difficult to detect with current comparative approaches (Pennell et al., 2015).

African cichlids (Ovalentaria, Cichliformes, Cichlidae, Pseudocrenilabrinae) provide a powerful system for investigating trait evolution dynamics. The subfamily is best known for the independent, adaptive radiations of the Great Rift Valley lakes Tanganyika, Malawi and Victoria, where remarkable species richness (McGee et al., 2020), morphological (Ronco et al., 2021; Malinsky et al., 2018) and behavioural diversity (Sommer-Trembo et al., 2024) has rapidly arisen. Seeding of the lake radiations occurred at different times during the diversification of Pseudocrenilabrinae: Lake Tanganyika radiation at approximately 12 million years old (Ronco et al., 2021), Malawi at 800 thousand years old (Malinsky et al., 2018) and Victoria (and rift lakes) at just 150 thousand years old (Meier et al., 2017), representing a series of independent evolutionary replicates or “*replays*” (Gould, 1989) unfolding over different temporal scales. Considered alongside older riverine lineages, this system offers a unique opportunity to compare radiations at different stages of divergence and to assess whether predicted macroevolutionary dynamics emerge consistently through time.

Vertebrae are the primary serial units of the vertebral column (Ford, 1937), the central axis of the vertebrate body plan. The number, type, and shape of vertebrae have been extensively modified during teleostean diversification (Ward and Brainerd, 2007; Baxter et al., 2022). Variation in vertebral column structure is closely linked to changes in body shape, particularly body elongation, a major axis of morphological diversification in multiple teleostean lineages (Claverie and Wainwright, 2014). The addition of vertebrae is a primary driver of body elongation (Ward and Brainerd, 2007; Mehta et al., 2010) and remarkable diversity in both total vertebral counts and body elongation has evolved across teleosts, including in African cichlids (Figure 1). Vertebral counts are determined during axial segmentation during somitogenesis (Morin-Kensicki et al., 2002), providing a direct link between developmental processes and adult morphology (Soul and Benson, 2017). Consequently, macroevolutionary shifts in vertebral counts offer not only a quantifiable proxy of body shape evolution, but also a tractable means of assessing the evolutionary lability of the developmental mechanism underpinning its evolution.

**Figure 1.**
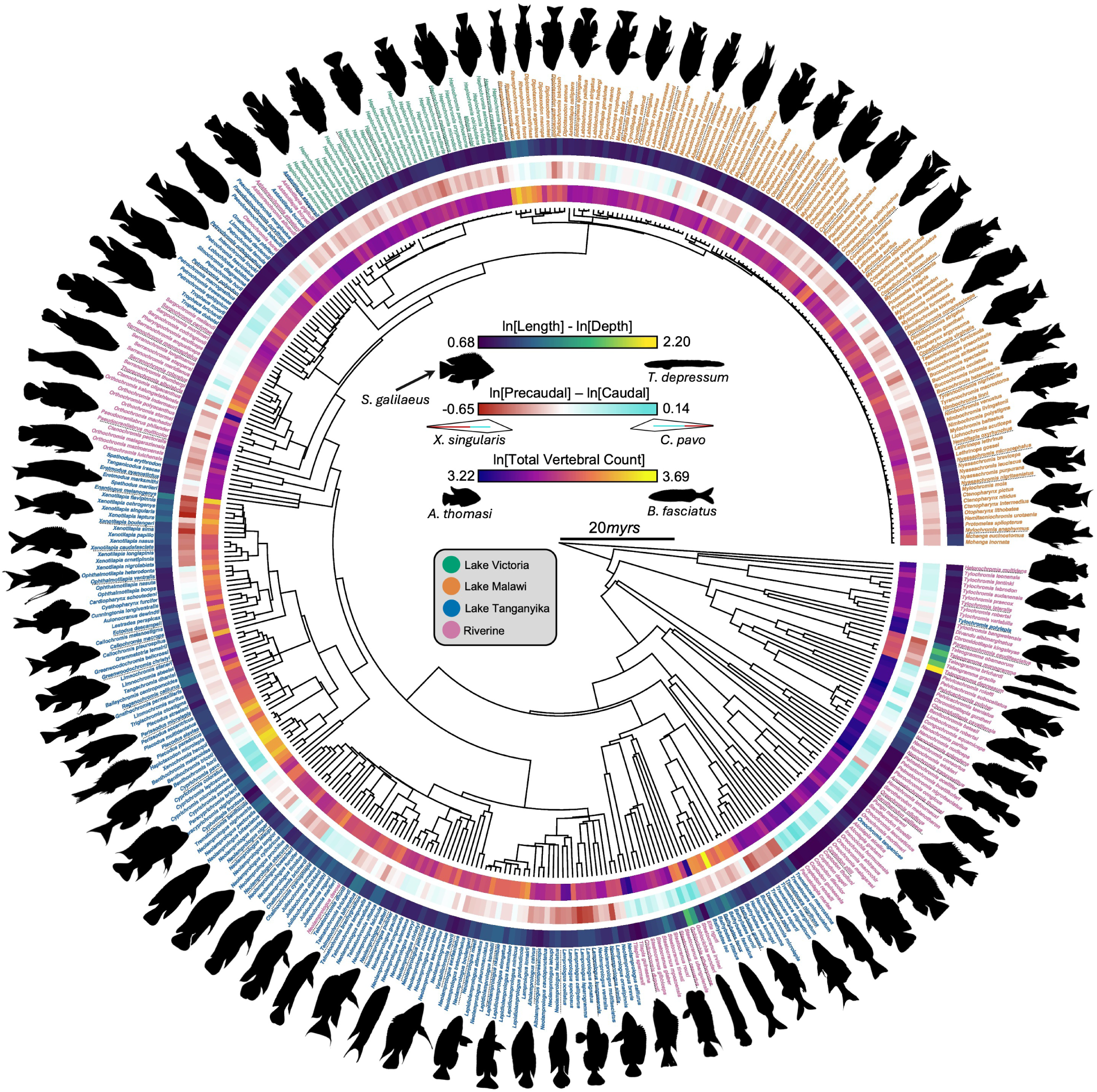
Vertebral counts and body shape have been modified during African cichlid diversification. Phylogeny from McGee et al. (2020) pruned to species present in the dataset (n=429 species). The respective ln[Total Count] (inner circle), ln[Precaudal] - ln[Caudal] (ratio of precaudal to caudal vertebrae, middle circle) and ln[Length] - ln[Depth] (body aspect ratio, outer circle) are indicated for each species as a heatmap (see legend, note that species shown represent the extremes). Silhouettes to demonstrate body shape diversity are shown for select species (dashed lines). Coloured tips indicate the water system occupancy of extant species (see legend). See Supplementary Materials for image acknowledgments. Abbreviations used in the figure: A. (*Anomalochromis*), B. (*Bathybates*), C. (*Cyprichromis*), S. (*Sarotherodon*), T. (*Teleogramma*), X. (*Xenotilapia*).

To date, no macroevolutionary study of the axial skeleton has been conducted in African cichlids, and comparative macroevolutionary analyses across their radiations remain scarce (Burress and Muñoz, 2023). Here, we compile a large comparative dataset of total, precaudal, and caudal vertebral counts, along with whole-body, cranial, and postcranial aspect ratios for 583 species of Pseudocrenilabrinae, representing all 26 currently recognised tribes (Astudillo-Clavijo et al., 2023), including both riverine lineages and major lacustrine radiations. Using this dataset, we test whether (i) evolutionary rates and variance in vertebral number scale predictably with radiation age, (ii) similar patterns of morphological disparity can arise from distinct underlying evolutionary processes, and (iii) macroevolutionary dynamics associated with adaptive radiation are more readily detectable when comparing radiations across temporal stages rather than within individual radiations.

By comparing Brownian motion (BM), Ornstein–Uhlenbeck (OU), and related models, we model ln[Total Count] evolution across the subfamily and within individual systems to assess rate variation through time and to test whether subfamily-level patterns are recapitulated within independent radiations. In doing so, we treat replicated radiations of African lake cichlids within Pseudocrenilabrinae as representing a common macroevolutionary process sampled at different time points. While this simplifying assumption may not fully capture ecological and evolutionary differences among systems (Malinsky et al., 2018; McGee et al., 2020; Ronco et al., 2021; Meier et al., 2017), it is heuristically useful for comparing evolutionary dynamics across radiations. This framework allows us to assess whether repeated radiations can be treated as comparable evolutionary time points, providing insight into how macroevolutionary patterns scale with time and why key signatures of adaptive radiation are inconsistently detected.

## METHODS AND MATERIALS

### Vertebral Count and Body Shape Measures

X-rays were collated from multiple sources for 4544 individuals, representing a total of 583 species (see Supplementary Methods). All x-rays have been deposited online (Bucklow et al., 2026). Images were landmarked in FIJI (Schindelin et al., 2012; Schneider et al., 2012) using predefined anatomical landmarks (Supplementary Figure 1). Vertebral centra were counted from anterior to posterior, including the urostyle, with fused centra inferred from neural spine counts. Vertebrae were classified as precaudal or caudal based on rib association and haemal spine morphology, with transitional forms assigned according to the presence of a haemal spine (and therefore haemal arch) ((Ford, 1937); see Supplementary Methods). Body shape was quantified using three aspect ratios (whole-body, anterior, and posterior) derived from landmark coordinates (Ronco et al., 2021). As scale bars were often absent, measurements were taken in pixels and expressed as ratios of log-transformed values. Specimens with incomplete anatomical data were excluded (n = 71).

### Phylogenetic Comparative Methods

All analyses were conducted in R (v4.2.0) (R Core Team, 2022). Phylogenetic analyses used the cichlid phylogeny constructed by McGee et al. (2020) as the tree is ultrametric, time-calibrated and includes all three of the major lacustrine radiations of African cichlids (Lake Tanganyika, Lake Malawi and Lake Victoria), as well as representatives from every currently recognised tribe in the African subfamily. After pruning the tree and subsetting our dataset, we were left with data for 429 species (131 genera) of Pseudocrenilabrinae, with representatives from all tribes within the subfamily (Astudillo-Clavijo et al., 2023). A discussion of the taxonomic considerations, including defining water system occupancy, can be found in the Supplementary Materials.

#### Phylogenetic Principal Component Analysis

We used phylogenetic principal component analysis (PPCA) to identify the primary the axes of variation and quantify occupied axial morphospace in our dataset (Revell, 2009). We input our count data (ln[Total Count], ln[Precaudal] and ln[Caudal]) and transformed body aspect ratios (anterior, posterior and whole) into the *phyl.pca* function in the R package *phytools* (v2.1.1) (Revell, 2024) with a BM-derived correlation matrix (Supplementary Table 1). We clustered species according to their respective lacustrine (Lake Tanganyika, Lake Malawi or Lake Victoria) or riverine system occupation (see Figure 1), their placement on the benthic-pelagic axis and for diet preference (piscivore versus non-piscivore). The 95% CI for group was considered to be the occupied axial morphospace for each respective group. To quantify overlap, we generated 100,000 random points within the lacustrine morphospace using *mvrnorm* (*MASS* v7.3.58.2) (Venables and Ripley, 2002). We then calculated each point’s Mahalanobis distance (*T*^2^) from the riverine centroid, classifying points as within the riverine morphospace if *T*^2^ *≤* 5.991, the 95% confidence threshold for a Chi-square (*χ*^2^) distribution with 2 degrees of freedom. Percentage overlap was defined as the proportion of lacustrine points within this boundary. A multivariate analysis of variance (MANOVA) was used to test for significant differences in occupied axial morphospace between groups. This was implemented using the *procD.lm* function in the R package *geomorph* (v4.0.6) (Baken et al., 2021), which performs MANOVA via residual randomisation, as provided by the *RRPP* package (Collyer and Adams, 2018). Pairwise comparisons were made using the *pairwise* function in *geomorph* (Baken et al., 2021).

#### Phylogenetic Linear Regression

Phylogenetic generalised least squares (PGLS) models were used to evaluate the evolutionary relationships between our univariate traits (Supplementary Table 2). Models were fit using the *pgls* function in the R package *caper* (v1.0.1). Trait variance-covariance matrices were transformed using Pagel’s *λ* (Pagel, 1997, 1999). To test whether the scaling relationship between vertebral count and body elongation varied among lineages, we fit a phylogenetic linear mixed model with total vertebral count as a fixed effect and species-specific random slopes whose covariance structure was defined by the phylogeny. Models were fit using the *pglmm* function in the R package *phyr* (v1.1.0) (Li et al., 2020). To test for significant differences in univariate group means, we used the *phylANOVA* function in the R package phytools (v2.1.1) (Revell, 2024), which implements the simulation-based phylogenetic ANOVA of Garland Jr et al. (1993). Each test was run with 100,000 simulations, and multiple testing was accounted for using a Holm–Bonferroni correction. To assess the relationship between total count and body aspect ratio within species, we fit within-species regressions (ordinary least squares) of ln[Total Count] on ln[Length] – ln[Depth] (body aspect ratio) for species with at least 10 individuals (176 species). To assess the overall subfamily pattern, we used a multivariate phylogenetic meta-analysis (Lajeunesse, 2009) using restricted maximum likelihood (REML) via the *rma.mv* function in the R package *metafor* (v4.8.0) (Viechtbauer, 2010). Species-specific regression slopes were treated as effect sizes and weighted by their respective sampling variances (i.e. the squared standard errors). Species were included as a random effect, allowing each to deviate from the overall mean slope. To account for shared evolutionary history, we included a phylogenetic covariance matrix constructed using the *vcv* function from the R package *ape* (v5.7.1) (Paradis and Schliep, 2019).

#### Ancestral Discrete Trait Estimation

We estimated the ancestral states of five of the seven univariate traits, including the total count (Supplementary Figure 9), inferring possible combinations of precaudal and caudal vertebrae from the ancestrally reconstructed precaudal:caudal ratio (ln[Precaudal]-ln[Caudal]) and ln[Total Count], using single-rate BM models in *fitContinuousMCMC* in the R package *geiger* (v2.0.10) (Pennell et al., 2014). Chain mixing and convergence was assessed in the R package *coda* (v0.19.4.1) (Plummer et al., 2006). For each univariate trait we combined the results from five independent Markov chains that ran for 2 million generations and discarded the first 10% as burn-in. Effective sample sizes were all greater than 250. We identified two extant species that closely resembled the reconstructed common ancestor, one of which most closely matched in axial morphology alone, and another that most closely matched in terms of axial morphology and in ancestrally-reconstructed discrete traits. For the former, we took the median (*Z*_0_) for each ancestrally reconstructed univariate trait and calculated the Euclidean distance between the ancestor and each extant species using the mean for each of the five reconstructed traits. For the inclusion of discrete traits, we calculated the Gower distance between the ancestral and extant species using the *daisy*function in the R package, *cluster* (v2.1.4) (Maechler et al., 2025). To quantify the similarity of these species with the common ancestor we also calculated similarity scores (1 *−D_min_/D_max_*), where *D_min_/D_max_*represents the ratio between the lowest (most similar) distance and highest (least similar) distance score within the extant species.

Discrete traits, including water system occupancy (Supplementary Figure 2), presence on the benthic-pelagic axis (Supplementary Figure 3) and piscivory (Supplementary Figure 4) were ancestrally reconstructed using the *fitDiscrete* function in the R package *geiger* (v2.0.10) (Pennell et al., 2014). Ecological data was mined from FishBase (Froese and Pauly, 2000). To allow more comparisons downstream, we pruned the tree to the minimal tree (i.e. species for which we had a complete ecological dataset), leaving us with a tree with 415 tips. We fit an equal rates (ER) model, where a single parameter governs all transition rates, an all-rates-different (ARD) model, where each transition has a unique rate parameter, and a symmetric (SYM) model, where forward and reverse transitions share the same parameter. Where only two groups were present (riverine versus lacustrine and piscivory versus non-piscivory), we omitted a SYM model fit, as a SYM model collapses to an ER model when the number of parameters (k) = 2 (Pagel, 1994; Lewis, 2001). As determined by comparison of the AICc for each model fit, for all but the haplochromines (Pseudocrenilabrini; (Oliver, 2024)), versus Lake Tanganyika endemics and basal riverine lineages (Supplementary Figure 5), which reported an ER model as the best fit, an ARD model was the best model fit. To account for uncertainty in the model fit, we simulated the best fitting model 1000 times on the minimal tree in *make.simmap* also in *geiger* (v2.0.10) (Pennell et al., 2014). For inference of qualitative traits at the common ancestor of African cichlids, we additionally included brooding behaviour and repeated model fitting for piscivory and benthic–pelagic occupancy using branch lengths scaled to one (Supplementary Figure 6).

#### Modelling ln[Total Count] Evolution

To investigate the evolutionary dynamics of ln[Total Count] across Pseudocrenilabrinae, we tested four non-mutually exclusive hypotheses. First, we evaluated whether variance in ln[Total Count] primarily reflects differences in stochastic evolutionary rates between riverine and lacustrine environments. Second, whether ln[Total Count] variance is primarily a function of radiation age, consistent with time-dependent divergence occurring either at constant rates or under lineage-specific stochastic rates. Third, given the exceptionally high haplochromine diversification rate (McGee et al., 2020), which includes the Lake Malawi and Lake Victoria radiations and an additional Tanganyikan radiation (Tropheina) (Ronco et al., 2020, 2021), we asked whether ln[Total Count] evolution differs within Pseudocrenilabrini (haplochromines) relative to the remainder of the Lake Tanganyika radiation and non-haplochromine riverine lineages, Finally, we examined whether ln[Total Count] evolution is best explained by ecological regime, as pelagic and piscivorous cichlids have independently evolved elongate bodies supported by increased vertebral counts (Stiassny, 1981; Vranken et al., 2023) (this study, Supplementary Figure 7).

Although total vertebral count is a discrete trait, intraspecific variation within species permits mean vertebral counts to be treated as continuous for comparative analyses (Supplementary Figure 8). Therefore, we employed a series of comparative analyses, comparing the fit of Brownian motion (BM) and Ornstein-Uhlenbeck (OU) models of continuous trait evolution on regimes that we ancestrally reconstructed on the phylogeny. The fit of multiple BM and OU models was tested amongst our regimes in *OUwie* (Beaulieu and O’Meara, 2022) on a subset of 100 trees from our discrete mapping to account for variability in the model fits. This included a single rate (*σ*^2^) and multi-rate BM model (BMS), in which each regime is permitted its own *σ*^2^, as well as the comparison of multiple OU models, where the attraction (*α*), rate (*σ*^2^) and trait optima (*θ*) parameters are constant or variable between regimes. Median AICc values were used to select the best fitting model within a regime and the best overall fitting model was selected using the AICc weights (see Table 1).

**Table 1.**
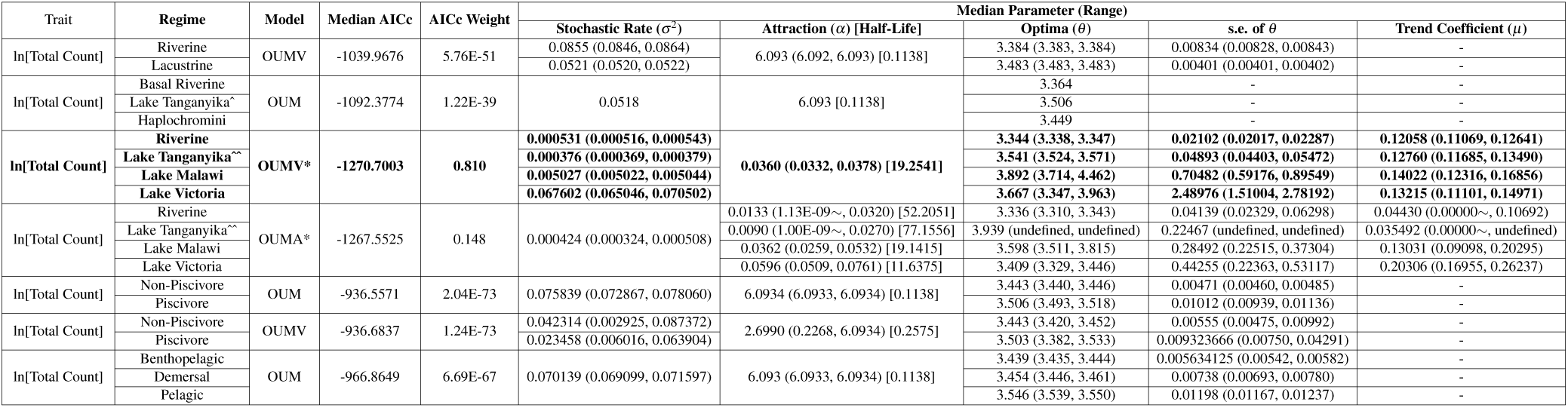
The parameters of the best fitting models per regime for ln[Total Count] evolution. Results are drawn from single and multi-regime Brownian motion (BM) and Ornstein-Uhlenbeck (OU) models, only OU models fit best for each regime. Parameters were estimated from 100 trees with ancestral estimates of the indicated regime (see Methodology). The median Akaike’s information criteria for finite sample sizes (AICc) and the Median AICc weight (global between regimes) is indicated for each model. In all OUM-class models (OUM, OUMA, OUMV), the trait optimum of ln[Total Count] (*θ*) was allowed to vary among regimes. In OUM, both the strength of stabilising selection (*α*) and the stochastic rate of evolution (*σ*^2^) were shared across regimes; in OUMA, *α* varied among regimes while *σ*^2^ was shared; and in OUMV, *σ*^2^ varied among regimes while *α* was shared. The phylogenetic half-life (*ln*(2)*/α*) is time taken in millions of years for half of the phylogenetic covariance to be erased between sister taxa. *When *α ∼* 0, the OU model behaves similarly to a BM model with a linear directional trend coefficient (*µ* = *αθ*) which represents the change in ln[Total Count] per million years. The model with the highest support amongst all model fits is shown in bold. No uncertainty is shown for Basal Riverine, Lake Tanganyika, or Pseudocrenilabrini (haplochromines) because regime assignments did not vary across the stochastic character maps. ^Does not include Tropheina. ^^Includes Tropheina.

We also assessed the fit of water-system–specific time-dependent models on pruned subtrees using the early-burst (EB; also referred to as ACDC) using *fitContinuous* in the R package (Pennell et al., 2014). In the ACDC model, the instantaneous evolutionary rate (*σ*^2^, the variance) changes exponentially through time according to 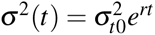, where 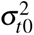 is the rate at the root and *r* is a rate-scaling parameter. Values of *r >* 0 indicate an increasing rate of variance accumulation through time, whereas *r <* 0 indicate a decreasing rate (effectively an early-burst dynamic). When *r ≈* 0, the model approximates a constant-rate Brownian motion (BM) process (Blomberg et al., 2003; Harmon et al., 2010). Relative model support was evaluated using AICc weights (Supplementary Table 3) of a comparison of an ACDC model against a single-rate BM model, an OU model, and a white-noise model, the latter assuming each species trait is an independent sample from a Gaussian distribution estimated by the respective the sample mean, *µ* and variance, *σ*^2^ (effectively independent of phylogeny). For these analyses we considered two riverine clades, defined by the presence or absence of haplochromines, to account for the possible lacustrine origins of genera such as *Serranochromis* (Joyce et al., 2005). Parameter estimates for models receiving non-negligible support are presented in Supplementary Table 4.

#### Ancestral Continuous Trait Estimation

We estimated the ancestral states of five of the seven univariate traits, including the total count (Supplementary Figure 9), inferring possible combinations of precaudal and caudal vertebrae from the ancestrally reconstructed precaudal:caudal ratio (ln[Precaudal]-ln[Caudal]) and ln[Total Count], using single-rate BM models in *fitContinuousMCMC* in the R package *geiger* (v2.0.10) (Pennell et al., 2014). Chain mixing and convergence was assessed in the R package *coda* (v0.19.4.1) (Plummer et al., 2006). For each univariate trait we combined the results from five independent Markov chains that ran for 2 million generations and discarded the first 10% as burn-in. Effective sample sizes were all greater than 250. We identified two extant species that closely resembled the reconstructed common ancestor, one of which most closely matched in axial morphology alone, and another that most closely matched in terms of axial morphology and in ancestrally-reconstructed discrete traits. For the former, we took the median (*Z*_0_) for each ancestrally reconstructed univariate trait and calculated the Euclidean distance between the ancestor and each extant species using the mean for each of the five reconstructed traits. For the inclusion of discrete traits, we calculated the Gower distance between the ancestral and extant species using the *daisy* function in the R package, *cluster* (v2.1.4) (Maechler et al., 2025). To quantify the similarity of these species with the common ancestor we also calculated similarity scores (1 *−D_min_/D_max_*), where *D_min_/D_max_* represents the ratio between the lowest (most similar) distance and highest (least similar) distance score within the extant species.

### Data and Code Availability

All code used in the analysis has been deposited on GitHub and can be accessed here. All radiographs, count and body shape data has been deposited on Zenodo (Bucklow et al., 2026), where it can be freely downloaded, shared and used in further studies.

## RESULTS

### Vertebral addition is important but not the only driver of body elongation in African cichlids

Axial morphology is highly variable across Pseudocrenilabrinae, encompassing several-fold differences in body elongation and marked shifts in vertebral number and precaudal-caudal proportions (Figure 1). Across the subfamily, whole-body aspect ratio scales positively and non-linearly with total vertebral count (log–log, *β*_1_ *>* 1; Figure 2A; PGLS: *R*^2^ = 0.250, *β*_1_ = 1.511, *λ* = 0.918, *p <* 0.0001, d.f. = 427), and this relationship is highly conserved across the phylogeny (phylogenetic slope variance = 2.89 *×* 10*^−^*^4^; residual variance = 3.38 *×* 10*^−^*^6^), consistent with patterns observed across teleosts (Ward and Brainerd, 2007; Mehta et al., 2010; Ward and Mehta, 2010). However, this macroevolutionary scaling is not recapitulated within species. Intraspecific slopes are highly heterogeneous and, on average, not significantly different from zero, indicating that individuals with more vertebrae are not consistently more elongate. This decoupling likely reflects strong constraint on vertebral number within species, which typically varies by no more than *±*1 vertebra (Supplementary Figure 7). Consequently, the positive relationship between vertebral count and elongation arises primarily through divergence among lineages over evolutionary time, rather than through variation within species.

**Figure 2.**
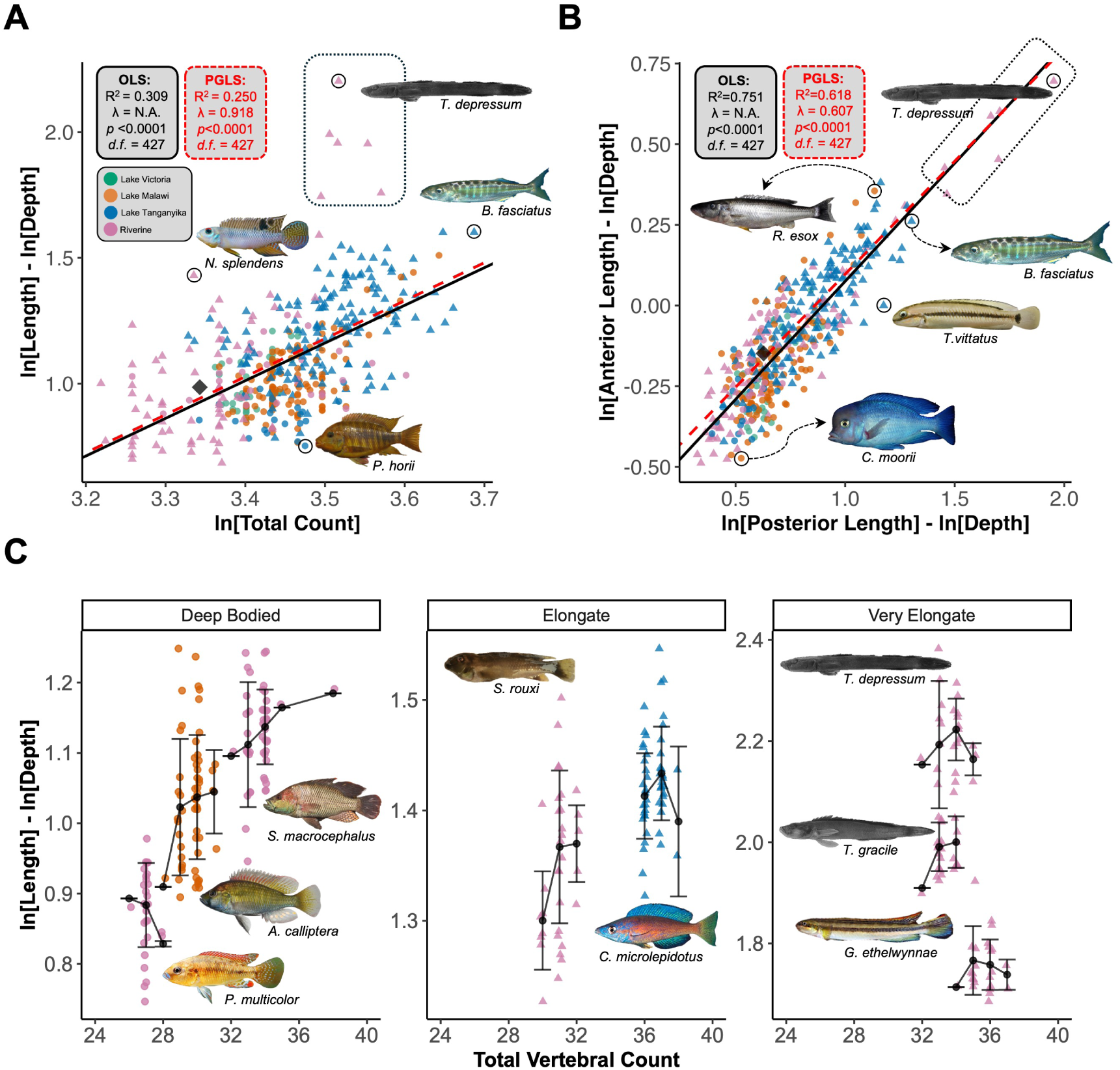
Vertebral addition is important but not the only driver of body elongation in African cichlids. (A) Relationships between whole body and ln[Total Count], and (B) anterior versus post-cranial aspect ratios, showing OLS (black) and PGLS (red, dashed) regressions (Supplementary Table 2). (C) Body aspect ratios plotted against vertebral count for selected species with more than 25 individuals. Panels are faceted by elongation category. Black circles represent the mean body aspect ratio for each vertebral count, with vertical bars indicating the standard deviation. See Supplementary Materials for image credits.

Surprisingly, vertebral counts account for relatively little of the variation in body elongation (*R*^2^ = 0.25). *Gobiocichla* and *Teleogramma*, for example, are highly elongate genera with considerably fewer vertebrae than predicted by the macroevolutionary scaling relationship (Figure 2B, dotted black box). Despite this, both *Teleogramma* and *Gobiocichla* possess elongate crania and post-cranial bodies consistent with the subfamily pattern (Figure 2B; *R*^2^ = 0.618, *β*_1_ = 0.688, *λ* = 0.738, *p <* 0.0001, *d.f.* = 427). In contrast, cranial elongation shows an even weaker association with total vertebral count (Supplementary Table 2, *R*^2^ = 0.085). Notably, both patterns are consistent with those observed in extremely slender elopomorphs (Mehta et al., 2010), suggesting these trends may be widespread among teleosts. Nonetheless, vertebral count only modestly explains body elongation evolution across the subfamily, within-species variation is minimal, and cranial elongation is largely independent of vertebral count evolution.

### Axial morphospace is primarily defined by body elongation and vertebral column regionalisation

Both total vertebral count and whole-body aspect ratio load strongly onto the dominant axis of axial morphospace across Pseudocrenilabrinae, indicating that the macroevolutionary association between vertebral number and body elongation strongly structures overall axial morphological diversity, even if variation in body elongation is not solely explained by vertebral counts (Figure 2A). The first principal component (PC1) explains 66.40% of the variation in axial morphospace within the subfamily, representing a linear combination of all seven traits (Figure 3A; Supplementary Table 1), including both total vertebral counts and body aspect ratio. Species scoring positively on PC1 have higher total, precaudal, and caudal vertebral counts, along with relatively elongate heads and postcranial bodies.

**Figure 3.**
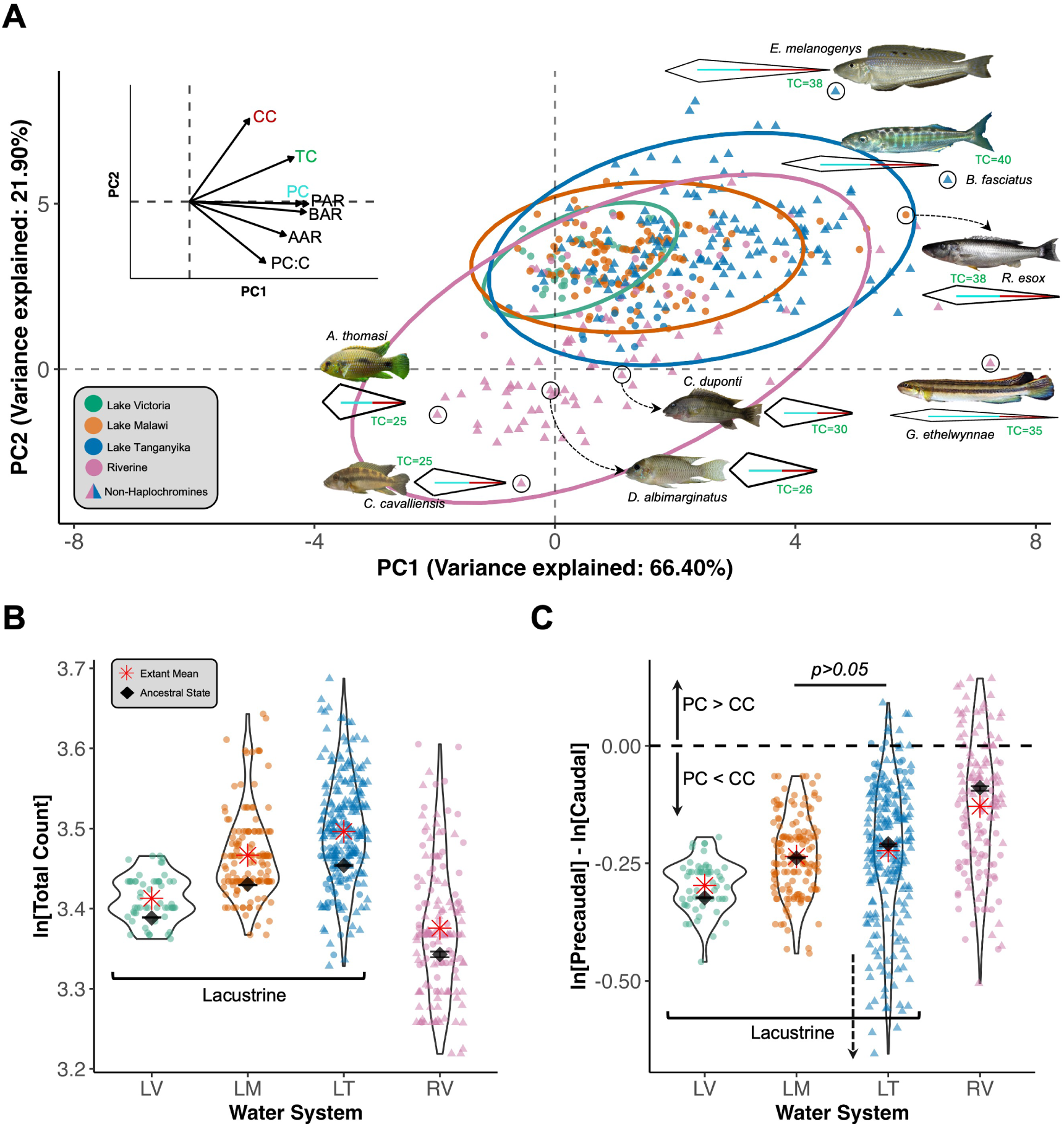
Lake taxa occupy a distinct subregion of riverine axial morphospace. (A) Phylogenetic PCA of axial traits showing PC1 versus PC2, with the proportion of variance explained by each axis indicated. Trait loadings are shown in the upper left. Species at the extremes of PC space and taxa closely resembling the inferred ancestral phenotype of Pseudocrenilabrinae are highlighted; modal total vertebral counts are shown in green. Ellipses represent 95% confidence intervals for each water system. Distributions of ln[Total Count] (B) and precaudal–caudal ratios (C) by water system, with crosses indicating means; all mean differences are significant (p *<* 0.001) except between Lake Tanganyika and Lake Malawi for the precaudal–caudal ratio. Ancestral estimates for both traits are indicated as black diamonds which represent the mean *±* 95%CI of estimated values at the respective ancestral nodes for each water system. Note that the riverine ancestor is the ancestor of all African cichlids (i.e. the root node). Abbreviations used: AAR (anterior aspect ratio, cranial), BAR (whole-body aspect ratio), CC (caudal vertebral count), PAR (posterior aspect ratio, postcranial), PC (precaudal vertebral count), PC:C (ratio of precaudal to caudal vertebrae), TC (total vertebral count). See Supplementary Materials for image credits.

While PC1 captures the primary axis of elongation, variation along the second principal component (PC2) reflects a complementary pattern of vertebral column regionalisation. PC2 explains 21.90% of axial morphospace variation, with positive values corresponding to high caudal vertebral counts and low precaudal:caudal ratios, whereas species with more negative PC2 values have lower absolute caudal vertebral counts (Figure 3A). Together, PC1 and PC2 account for 88.3% of total axial variation, indicating that a considerable proportion of axial morphological diversification in African cichlids occurs within a shared, low-dimensional morphospace defined primarily by body elongation, driven in part, but not solely by, vertebral addition, and vertebral column regionalisation.

### Lake taxa occupy a distinct subregion of riverine axial morphospace

Riverine morphospace is substantially larger than lacustrine morphospace, indicating that the lake radiations have largely resampled axial variation already present among the older, riverine lineages (Figure 3A). Despite this, riverine and lacustrine taxa occupy significantly different regions of morphospace (MANOVA, PC1–PC4, *R*^2^ = 0.191, F(1,427) = 100.97, Z = 8.49, *p <*0.0001; Supplementary Figure 10). Riverine taxa are concentrated toward the negative extremes of PC1 and PC2, encompassing both small-bodied ‘dwarf’ cichlids (e.g. *Anomalochromis thomasi*, *Chromidotilapia cavalliensis*) and large, deep-bodied lineages with relatively few vertebrae (e.g. Oreochromini, Coptodonini, Tilapiini, Pelmatolapiini; (Astudillo-Clavijo et al., 2023)). In contrast, lacustrine taxa occupy higher values along both axes, with extremes represented by species such as *Enantiopus melanogenys* and pelagic piscivores including *Bathybates fasciatus* and *Rhamphochromis esox*. Consistent with these distributions, 90.43% of lacustrine morphospace is occupied by riverine taxa, whereas 49.96% of riverine morphospace is unique to riverine taxa, reinforcing that lacustrine radiations expand within, rather than beyond, the broader riverine morphospace. Excluding haplochromines (Pseudocrenilabrini) increases the separation between riverine and lacustrine (now just Lake Tanganyika) assemblages, with lacustrine taxa occupying a more discrete subregion of axial morphospace (MANOVA, PC1–PC4, *R*^2^ = 0.362, F(1,224) = 126.82, Z = 8.137, *p <*0.00001; Supplementary Figure 11). Nonetheless, overlap remains substantial: 57.32% of lacustrine morphospace is still occupied by riverine taxa, indicating that even in the absence of haplochromines, Lake Tanganyikan cichlids have diversified within pre-existing riverine axial variation. Riverine species are, on average, deeper-bodied and possess fewer caudal vertebrae than lacustrine species (MANOVA, *d* = 2.742, Z = 8.218, *p <*0.0001), whereas lacustrine taxa exhibit higher total vertebral counts and lower precaudal:caudal ratios (Figures 3B, C). Ancestral state estimates indicate that the common ancestor of African cichlids had a low vertebral count (28.22 [28.19, 28.39]) and was likely deep-bodied (Figure 2A; (Oliver, 2024)). Ancestral estimates further support a shallow, benthopelagic riverine habitat, substrate brooding, and a non-piscivorous diet (Supplementary Figure 6). This inferred ancestral condition occupies a region of axial morphospace low in both PC1 and PC2, consistent with extant riverine taxa such as *Divandu albimarginatus* and *Chilochromis duponti* (Figure 3A), indicating that the ancestral phenotype falls within a portion of morphospace still unique to riverine lineages.

### Parallel increases in vertebral number and body elongation occur across lacustrine radiations

Lacustrine radiations occupy nested regions of morphospace (Figure 3A). Older radiations have explored a broader portion of axial morphospace, and their occupied regions encompass those of younger radiations. Ancestral vertebral counts for the lineages seeding Lakes Tanganyika (31.12), Malawi (31.63), and Victoria (30.05) are within the expected intraspecific range (Supplementary Figure 7) and lower than their respective extant lake means (Figure 3B), suggesting the repeated parallel evolution of elongate bodies with increased vertebral counts. In contrast, ancestral estimates of the precaudal–caudal ratio differ little from extant means (Figure 3C), suggesting that lake seeders already possessed relatively low ratios. Therefore, most lake morphological change has focused on increasing total vertebral counts (precaudal and caudal) and body elongation (along PC1), rather than the preferential addition of caudal vertebrae. Consequently, the skewed ratio distribution observed in Lake Tanganyika (Figure 3C, black arrow) may reflect lineage-specific caudal vertebral addition in taxa occupying high PC2 morphospace, such as the Ectodini and pelagic piscivores such as those belonging to *Bathybates* (see Supplementary Figure 7) that coincidentally occupy a portion of the axial morphospace unique to lacustrine taxa (Figure 3A).

### Differences in morphospace occupation emerge from water-system–specific stochastic rates and contrasting trait optima

Consistent with greater axial morphospace occupation in riverine taxa relative to lacustrine lineages, residual variance in ln[Total Count] is significantly higher in extant riverine species (Levene’s test: *F* = 10.747, *d.f.* = 427, *p* = 0.00113) and nearly twice that observed in lacustrine species (variance ratio = 1.78). However, total vertebral count disparity across the subfamily cannot be explained by adaptation to lacustrine versus riverine environments alone (AICc weight *≈* 0), haplochromine-specific dynamics (AICc weight *≈* 0), or ecological axes such as benthic–pelagic position or piscivory (AICc weights *≈* 0; Table 1), despite repeated evolution of increased vertebral counts and elongate bodies in pelagic and piscivorous lineages (Supplementary Figure 7; (Stiassny, 1981; Vranken et al., 2023)).

Of the models tested, ln[Total Count] evolution is best explained by water-system–specific stochastic rates (*σ*^2^) and contrasting trait optima (*θ*), with a shared strength of attraction toward those optima (median *α*; OUMV model, AICc weight = 0.810; Table 1). Under this model, riverine lineages were attracted toward a lower trait optimum (median *θ* = 3.344) than any of the lake radiations (3.541–3.892), consistent with their occupation of axial morphospace at lower PC1 and PC2 values (Figure 3A) and their lower mean ln[Total Count] (Fig. 3B). Riverine lineages date to the origin of Pseudocrenilabrinae (64–58 Ma; (Friedman et al., 2013; Matschiner et al., 2020)), spanning approximately three phylogenetic half-lives. Accordingly, ln[Total Count] is expected to have approached its stationary variance in riverine lineages but not in the lakes. The greater variance observed in riverine taxa therefore likely reflects their longer evolutionary history rather than weaker constraint.

### Axial disparity in lacustrine radiations is limited by lack of time to diversify

Lake occupation of axial morphospace and variance in ln[Total Count] correlate with radiation age (Figure 3A, B). In contrast to riverine lineages, the lake radiations are substantially younger than a single phylogenetic half-life (19.25 Myr versus *<*12 Myr; (Ronco et al., 2021)). Consequently, stochastic rates primarily reflect differences in the tempo of variance accumulation, as opposed to fluctuation around the trait optima. Rates of ln[Total Count] evolution (*σ*^2^) inversely correlate with radiation age, with the youngest radiations exhibiting the most rapid variance accumulation. Median net rates are the lowest in Lake Tanganyika (0.000376 ln[Total Count] Myr*^−^*^1^), an order of magnitude (13x) faster in Lake Malawi (0.005027), and faster still (179x) in Lake Victoria (0.067602; Table 1). Therefore, younger systems evolve more rapidly but have had insufficient time to accumulate substantial variance, whereas Lake Tanganyika has accumulated extensive variance over a longer time despite its average slower rate. Therefore, differences in axial morphospace occupation between the lake radiations has ultimately been limited by the time over which they have existed.

### Phylogenetic structure emerges across radiations through time

Model support within individual lakes, however, suggests differences in evolutionary mode, or that different processes of phenotypic diversification are important through the timescale of adaptive radiations. In Lake Victoria, strongest support for a white-noise model (AICc weight = 0.791; Supplementary Table 3) implies that phenotypic diversification is phylogenetically unstructured on short timescales. In the older Lake Malawi, equal support for OU and ACDC models (AICc weights = 0.500; Supplementary Table 3) suggests that both stabilising dynamics and exponentially accelerating variance accumulation (r = 2.174, *σ*^2^ = 3.67 *×* 10*^−^*^5^; Supplementary Table 4) can equally account for the observed variance structure. In contrast, in Lake Tanganyika uniform Brownian accumulation best explains ln[Total Count] evolution. We found equivalent support for a BM (AICc weight = 0.397), OU (AICc weight = 0.302), and very weakly accelerating (r=0.037, Supplementary Table 4), ACDC model (AICc weight = 0.302; Supplementary Table 3), with the latter essentially collapsing to a constant-rate BM model, indicating a high degree of phylogenetic structure in the older radiations. Notably, similar patterns were also observed in riverine lineages excluding haplochromines (compare AICc weights with Lake Tanganyika, Supplementary Table 3), with estimated Brownian motion (*σ*^2^ = 0.000204, Supplementary Table 4) rates closely matching those of Lake Tanganyika (*σ*^2^ = 0.000296, Supplementary Table 4). Therefore, over longer evolutionary timescales, variance accumulation may converge on similar net rates across systems. Taken together, these results indicate that similar variance patterns can arise from distinct evolutionary dynamics, reflecting a transition from phylogenetically unstructured to increasingly structured trait evolution across radiations (Figure 4).

**Figure 4.**
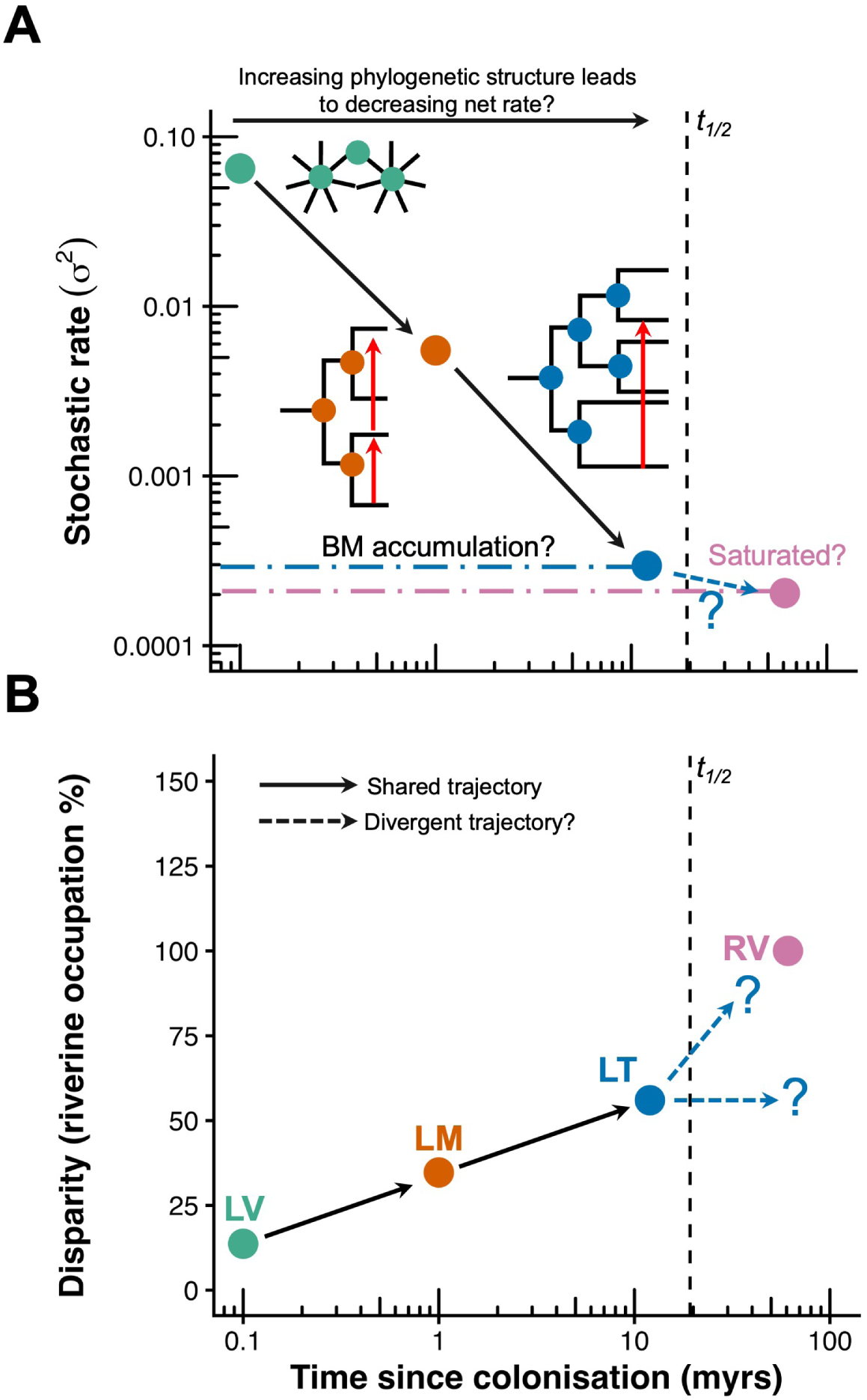
Increasing phylogenetic structure is associated with declining net evolutionary rates across replicated adaptive radiations. **(A)** Conceptual and empirical summary of stochastic rates of ln[Total Count] evolution across radiations of different ages. Younger systems exhibit higher net rates and reduced phylogenetic structure, whereas older radiations show lower rates and increased phylogenetic structuring, consistent with a temporal shift in underlying evolutionary dynamics. Schematic overlays illustrate one possible interpretation of this transition, in which early diversification is characterised by high reticulation (e.g. hybridisation or incomplete lineage sorting) that diminishes through time. Riverine lineages may have followed a similar trajectory (dashed line), but constant-rate (BM-like) accumulation cannot be excluded for these lineages and for Lake Tanganyika (dot-dash line). **(B)** Axial morphospace occupation through time, illustrating that younger radiations occupy a smaller proportion of total disparity, consistent with time-limited expansion. Arrows indicate hypothesised trajectories of morphospace exploration across radiations; however, these trajectories may be constrained in some systems (e.g. Lake Victoria and associated lakes) by ecological limits such as reduced depths.

## DISCUSSION

Adaptive radiations provide a natural framework for testing whether macroevolutionary dynamics are predictable across replicated evolutionary events. Pseudocrenilabrinae originated from a relatively deep-bodied, benthopelagic riverine ancestor with few vertebrae approximately 64–58 mya (Friedman et al., 2013; McGee et al., 2020; Matschiner et al., 2020). Since then, repeated transitions into lacustrine environments have generated multiple independent radiations characterised by rapid ecological and morphological diversification (Salzburger et al., 2005). Across these radiations, we find that lineages repeatedly explored the same regions of axial morphospace, indicating that diversification is structured by a combination of ancestral morphology and evolutionary rates. However, the total extent of morphospace occupation remains low, consistent with short divergence times limiting the accumulation of morphological variance of younger radiations. Therefore, our results imply that adaptive radiations follow predictable but temporally bounded trajectories, in which evolutionary outcomes are both repeatable in direction and limited in extent.

Although stochastic rates of evolution temporally scale such that younger clades tend to exhibit higher net rates of trait evolution (Gingerich, 1983), as in the case for ln[Total Count] evolution (Figure 4), rate disparity magnitude between riverine and lacustrine lineages matches that previously reported for craniofacial traits (Burress and Muñoz, 2023). Therefore, high rates in Lakes Victoria and Malawi are due to their relatively recent lake colonisations and unlikely to be a statistical artefact of clade age. Rapid niche diversification, ecological opportunity, and reduced initial competition following colonisation of lacustrine environments likely facilitated accelerated exploration of axial morphospace. These elevated rates are also consistent with theoretical expectations from adaptive radiation models, in which early phases of diversification are characterised by rapid trait evolution as lineages expand into unexploited niches (Simpson, 1953; Schluter, 2000; Stroud and Losos, 2016).

We did not detect evidence of early burst (EB) dynamics within any individual system, including Lake Tanganyika (Supplementary Table 3), despite previous evidence for EB in body shape evolution (Ronco et al., 2021). This finding adds evidence that EB signals are rarely recovered in individual clades (Harmon et al., 2010). However, this may reflect limited time depth, low statistical power, or challenges in distinguishing EB from alternative models (Slater and Friscia, 2019). Across these systems, we observe a consistent pattern of declining net rates through time (OUMV, Table 1) and younger sub clades exhibiting less variation than older, more inclusive clades (Figure 3B), both of which are hallmark expectations of EB models (Harmon et al., 2003, 2010). However, elevated rates over short timescales do not necessarily accumulate into long-term divergence (Uyeda et al., 2011), such that the high net rates observed in younger radiations may partly reflect transient, time-limited fluctuations rather than the early phase of an early-burst process. Nonetheless, this raises the possibility that the apparent rarity of early-burst dynamics in comparative analyses reflects the limited temporal scope of individual radiations and the restricted temporal scales accessible in contemporary datasets (Harmon et al., 2010; Slater and Friscia, 2019), rather than their true absence, although such dynamics may alternatively be genuinely rare or only detectable across replicated radiations.

Strong support for a white-noise model for Lake Victoria is particularly notable and suggests that rapid phenotypic diversification on short timescales lacks phylogenetic structure. This pattern may arise as a statistical consequence of extremely shallow divergence times and relatively few taxa (see Supplementary Figure 2), conditions under which phylogenetic covariance is difficult to detect, and support for comparative models becomes weakly identifiable (Butler and King, 2004). However, it may also reflect temporal variation in the processes underlying phenotypic diversification. The Lake Victoria radiation (at least within Lake Victoria) is frequently described as a hybrid swarm, with substantial reticulation among lineages (Meier et al., 2017, 2023). Such reticulate evolutionary histories erode tree-like phylogenetic structure and reduce detectable phylogenetic signal, potentially contributing to the observed support for a white-noise process. Under these circumstances, bifurcating tree-based comparative methods may inadequately capture covariance structure, necessitating network-based phylogenetic comparative frameworks (Bastide et al., 2018). Nonetheless, our results suggest that very early diversification may be dominated by microevolutionary processes that rapidly generate disparity. In older radiations, net rates are lower and phenotypic variation is phylogenetically structured (e.g. BM in Lake Tanganyika, Supplementary Table 4) consistent with a transition from early rapid divergence to longer-term dynamics (Figure 4).

Riverine lineages occupy a broader axial morphospace than lacustrine lineages, encompassing almost all of lake morphospace while extending into regions characterised by low total vertebral counts and deeper bodies, consistent with prolonged evolution around a lower total count optimum distinct from that of the lakes (Table 1). Deep-bodied forms are commonly associated with riverine environments (Pavey et al., 2010; Mercer et al., 2020; Suriyampola et al., 2023) and may enhance manoeuvrability in relatively turbulent, structurally complex habitats (e.g. wood debris, rocks). Nonetheless, rivers clearly generate substantial axial disparity and this is further exemplified by predominantly riverine Neotropical cichlids (Subfamily: Cichlinae) (Sparks and Smith, 2004; Smith et al., 2008), which exhibit morphologies not observed in Pseudocrenilabrinae, including extremely deep-bodied *Symphysodon* and *Pterophyllum*, and highly elongate *Crenicichla* with large vertebral and precaudal counts (Varella et al., 2023).

Lakes, however, are not devoid of innovation: multiple lacustrine lineages occupy morphospace not represented in rivers (high PC1 and PC2; Figure 3A), including elongate Ectodini (e.g. *Enantiopus melanogenys*) and pelagic piscivores such as *Rhamphochromis* and *Bathybates*, which exhibit the highest vertebral counts in the subfamily (this study; (Stiassny, 1981; Oliver, 2024)). This lacustrine-specific region of morphospace is relatively limited, likely reflecting reduced time for variance accumulation rather than constrained innovation. Nonetheless, lake adaptation repeatedly drove elongation and increased vertebral counts, particularly along the benthic–pelagic axis and in piscivores (Supplementary Figure 7), despite weak explanatory power for ln[Total Count] evolution overall (Table 1). Because these niches are largely lacustrine (Supplementary Figure 2-Supplementary Figure 4), they may elevate ln[Total Count] optima (Table 1) driving repeated parallel elongation in the lakes.

The lack of positive scaling between intraspecific vertebral count variation and body aspect ratio (Figure 2C, Supplementary Figure 8) suggests that increases in body elongation are unlikely to have arisen through gradual selection on individuals with increased vertebral counts and low intraspecific variation in vertebral number (Supplementary Figure 8) may reflect strong developmental canalisation. Although environmental perturbations can influence vertebral counts (Fowler, 1970; Tibblin et al., 2016; Campbell et al., 2021), vertebral number is tightly constrained, likely reflecting the multiple roles of somites during embryogenesis (Morin-Kensicki et al., 2002; van Eeden et al., 1996; Kuan et al., 2004; Loring and Erickson, 1987), such that substantial deviations are rare and likely deleterious. Rapid increases in vertebral number may therefore occur through rare, large changes to segmentation dynamics, such as shifts in the segmentation clock, axis elongation, or the amount of tissue available for segmentation (Taylor et al., 2026), rather than through gradual accumulation.

Although incremental change is possible, the scale and rapidity of divergence observed in Lakes Victoria and Malawi, despite low per-generation mutation rates and short divergence times (Malinsky et al., 2018; Svardal et al., 2021), are difficult to reconcile with strictly gradual models. Strong developmental canalisation may therefore restrict evolution to a limited set of viable developmental trajectories (Alberch, 1989). Consistent with this, ancestral state estimates indicate that shifts toward piscivory generally precede increases in vertebral count and body elongation, suggesting that ecological transitions may create conditions under which relatively large, developmentally biased phenotypic shifts, similar to the “hopeful monsters” of Alberch, are favoured, without invoking single-generation macromutation (Gould, 1977). Empirical examples support this pattern: elongate piscivores have evolved rapidly in Lakes Victoria (Vranken et al., 2023) and Malawi (*Rhamphochromis*; this study), the latter in *<*350 kyr (Malinsky et al., 2018). Such dynamics could be consistent with pulsed models of evolution, for example Lévy-flight models, in which long periods of stasis are punctuated by infrequent, large shifts in trait space (Landis and Schraiber, 2017). This could provide a powerful framework for understanding how canalisation, ecological opportunity, and rare large-effect shifts interact to drive rapid morphological evolution.

Critically, our results highlight the importance of temporal scale in shaping inferences about macroevolutionary dynamics. Because most empirical studies focus on individual clades, they capture only partial snapshots of longer-term evolutionary processes. As a result, key signatures predicted by adaptive radiation theory, including early bursts, rate slowdowns, and shifts in phylogenetic signal, may be difficult to detect or may appear absent. By comparing replicated radiations across temporal stages, these dynamics can instead be recovered as emergent patterns across systems. This suggests that their apparent rarity may reflect limits of temporal sampling rather than true absence. Treating independent radiations as comparable evolutionary time points therefore provides a general framework for understanding how macroevolutionary dynamics emerge through time. In this context, African cichlids, with multiple independent radiations spanning a wide temporal range, offer a uniquely powerful system for linking microevolutionary processes to macroevolutionary outcomes. Integrating across these scales will be essential for understanding the evolutionary dynamics that have shaped this remarkable group.

## SUPPLEMENTARY FIGURES AND TABLES

**Supplementary Table 1.**
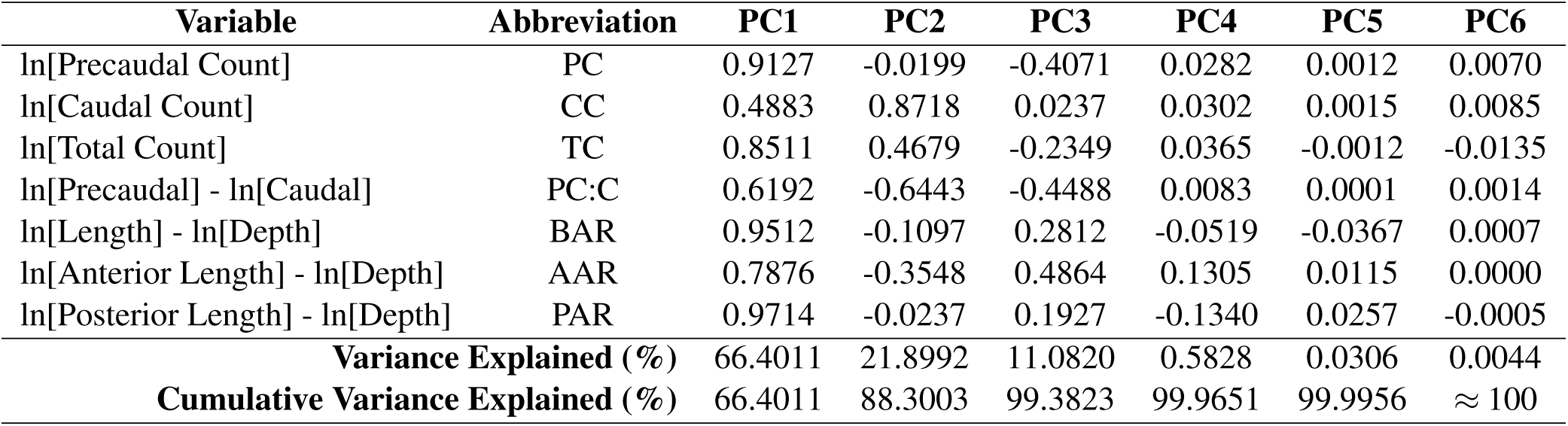
Phylogenetically-corrected principle component loadings for 429 species belonging to all tribes within Pseudocrenilabrinae. Abbreviations refer to those used in Figure 1 for each respective variable. Loadings for PC1 have been multiplied by −1 to make them positive (and consistent with Figure 3A). The contributions of PC7 are negligible and have been omitted.

**Supplementary Table 2.**
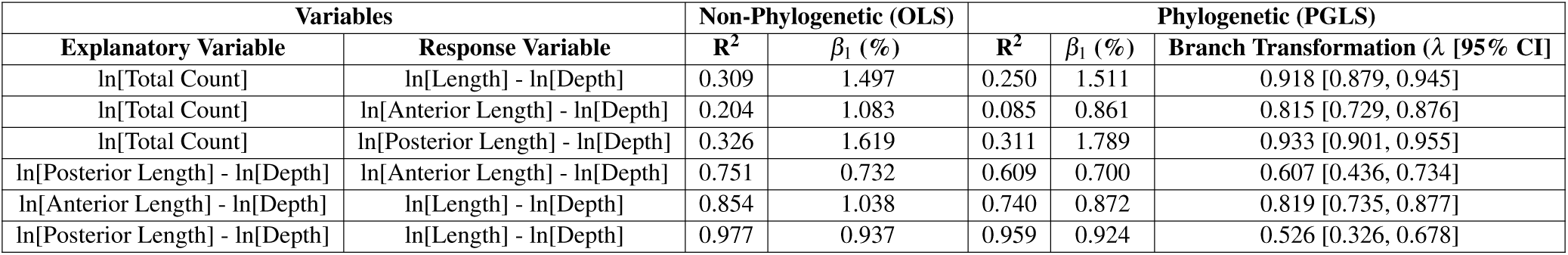
Univariate OLS and PGLS model fits are indicated for each relationship tested and based on data from 429 species retained on the McGee et al. (2020) phylogeny. Adjusted *R*^2^ values for each model fit are shown. Since all variables are natural log-transformed, the gradient/slope (*β*_1_) represents the *approximate* percentage change in the response variable for a 1% increase in the explanatory variable (elasticity). Maximum likelihood (ML) estimates for *λ* branch transformation are indicated along with their 95% confidence intervals. AIC weights for each PGLS model are indicated. Note that these weight values refer to the weight of the respective model fit for that relationship. All model fits are significant (*p <* 0.0001).

**Supplementary Table 3.** Support for all models fit to individual water systems. Models in bold have non-negligible support. Note that for multiple models have equal support for each water system. The number of parameters (k) and AICc values for each model fit are indicated. *Not comparable between water systems, only within. This is reflected in the ΔAICc and AICc weights which were calculated within the models tested for each water system.

| Model | Parameters (*k*) | AICc | ΔAICc* | AICc Weight* |
| --- | --- | --- | --- | --- |
| **Lake Victoria** |  |  |  |  |
| **OU** | **3** | **-94.030** | **2.657** | **0.209** |
| ACDC/EB | 3 | -51.622 | 45.065 | 0.000 |
| BM | 2 | -26.040 | 70.647 | 0.000 |
| **White** | **2** | **-96.687** | **0.000** | **0.791** |
| **Lake Malawi** |  |  |  |  |
| **OU** | **3** | **-392.004** | **0.000** | **0.500** |
| **ACDC/EB** | **3** | **-392.004** | **0.000** | **0.500** |
| BM | 2 | -376.434 | 15.569 | 0.000 |
| White | 2 | -378.873 | 13.131 | 0.001 |
| **Lake Tanganyika** |  |  |  |  |
| **OU** | **3** | **-595.071** | **0.550** | **0.302** |
| **ACDC/EB** | **3** | **-595.071** | **0.550** | **0.302** |
| **BM** | **2** | **-595.621** | **0.000** | **0.397** |
| White | 2 | -438.982 | 156.639 | 0.000 |
| **Riverine (no Haplochromines)** |  |  |  |  |
| **OU** | **3** | **-199.514** | **0.319** | **0.315** |
| **ACDC/EB** | **3** | **-199.514** | **0.319** | **0.315** |
| **BM** | **2** | **-199.833** | **0.000** | **0.370** |
| White | 2 | -155.861 | 43.972 | 0.000 |
| **Riverine (with Haplochromines)** |  |  |  |  |
| **OU** | **3** | **-269.134** | **0.000** | **0.499** |
| **ACDC/EB** | **3** | **-269.134** | **0.000** | **0.499** |
| BM | 2 | -256.904 | 12.230 | 0.001 |
| White | 2 | -210.090 | 59.044 | 0.000 |

**Supplementary Table 4.** Parameters of ln[Total Count] evolution models per water system with non-negligible support. For OU models, we report the attraction parameter (*α*), its half-life (ln(2)*/α*; Myr), and the stochastic rate (*σ*^2^; Myr*^−^*^1^). For EB (ACDC) models, the rate-scaling parameter (*r*) is shown: *r >* 0 indicates exponentially increasing variance 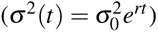, *r <* 0 a decreasing rate (early burst), and *r ≈* 0 approximates constant-rate BM. Root values (*Z*_0_) are provided. In the white-noise model, *Z*_0_ and *σ*^2^ correspond to the mean and variance of extant counts. ^Because the stationary variance of an OU process equals *σ*^2^*/*(2*α*), very large *α* values with correspondingly inflated *σ*^2^ effectively approximates a white noise model, or rather the extant mean and variance.

| Model | Attraction ($\alpha$) [Half-Life] | Scalar ($r$)* | Stochastic Rate ($\sigma^2$) | Root ($Z_0$) or Mean ($\mu$) |
| --- | --- | --- | --- | --- |
| **Lake Victoria** |  |  |  |  |
| White | — | — | 7.16e-04 | 3.413 |
| OU | 2262.845^ | — | 3.241^ | 3.413 |
| **Lake Malawi** |  |  |  |  |
| OU | 1.087 [1.155] | — | 7.18e-03 | 3.468 |
| ACDC/EB | — | 2.174 | 3.67e-05 | 3.468 |
| **Lake Tanganyika** |  |  |  |  |
| BM | — | — | 2.96e-04 | 3.456 |
| OU | 0.019 [36.481] | — | 3.34e-04 | 3.462 |
| ACDC/EB | — | 0.037 | 5.48e-05 | 3.462 |
| **Riverine (no Haplochromines)** |  |  |  |  |
| BM | — | — | 2.04e-04 | 3.333 |
| OU | 0.015 [45.210] | — | 2.67e-04 | 3.339 |
| ACDC/EB | — | 0.030 | 4.60e-05 | 3.339 |
| **Riverine (with Haplochromines)** |  |  |  |  |
| OU | 0.041 [16.906] | — | 5.75e-04 | 3.347 |
| ACDC/EB | — | 0.082 | 4.79e-06 | 3.347 |

**Supplementary Figure 1.**
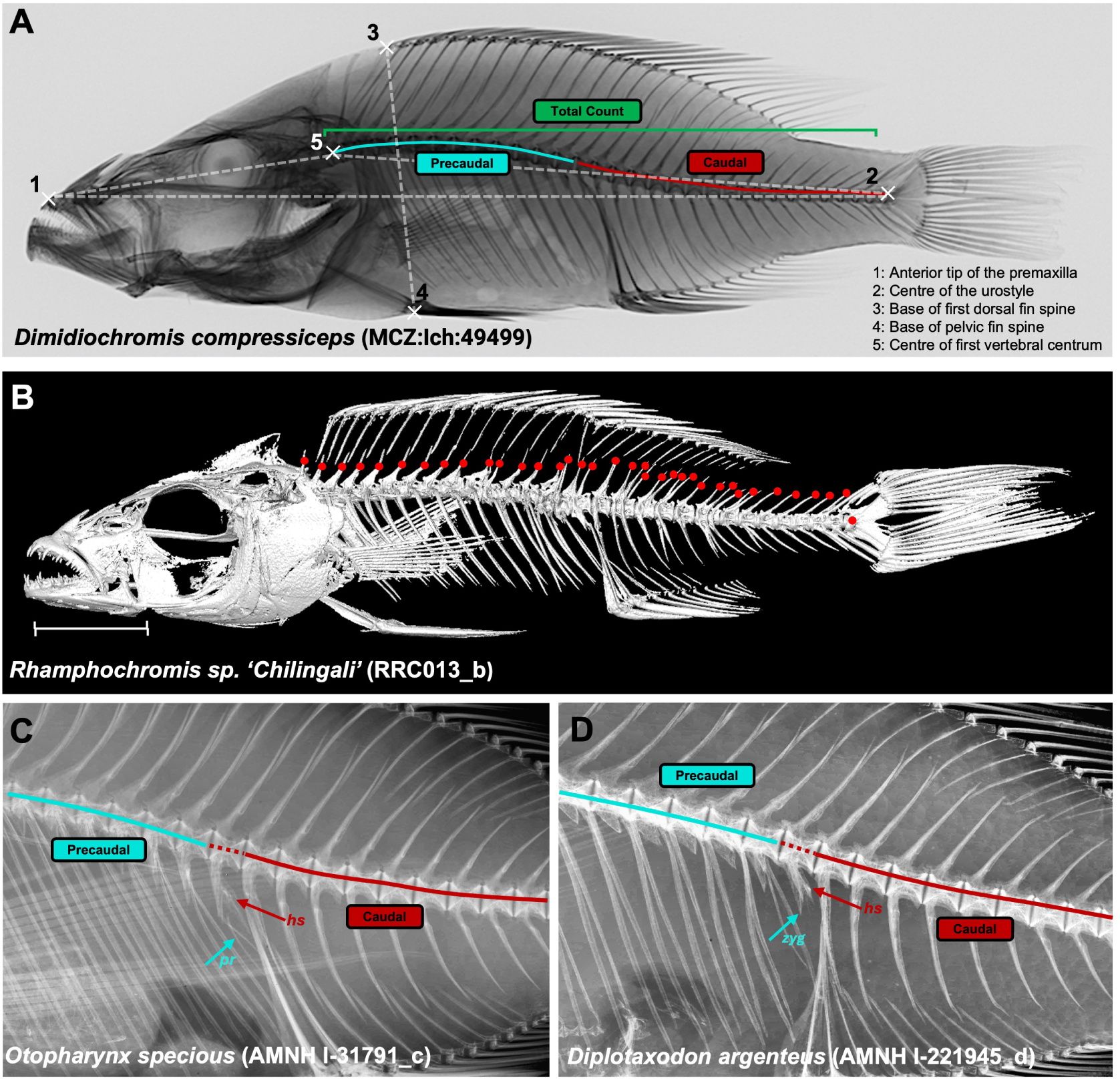
Measuring axial traits and body shape. (A) Landmarks used in the study, as previously published (Ronco et al., 2021). Dashed lines indicate straight lines used for the calculation of the three respective aspect ratios used in the study. Precaudal vertebrae are indicated in blue and the caudal vertebrae in red. The total vertebral count (green) is the summation of the precaudal and caudal vertebral counts. The specimen data record can be accessed here. (B) A specimen of *Rhamphochromis* sp. ‘Chilingali’ with a deformed vertebral column with multiple instances of fused vertebrae. Note that each fused centrum still retains a neural spine. Scale is 1cm. The threshold value used to generate the model was not high enough to resolve the epineural bone but it is indeed present in the specimen. (C) A transitional vertebra bearing both rudimentary pleural ribs (pr) (blue arrow) and a haemal spine (hs) and therefore a haemal arch. Specimen references are indicated for each image. (D) An example of a transitional vertebra that bears zygapophyses (zyg) resembling a precaudal vertebra (blue arrow) but has a rudimentary haemal spine (hs) and therefore a haemal arch. See Supplementary Materials for images matching these specimens.

**Supplementary Figure 2.**
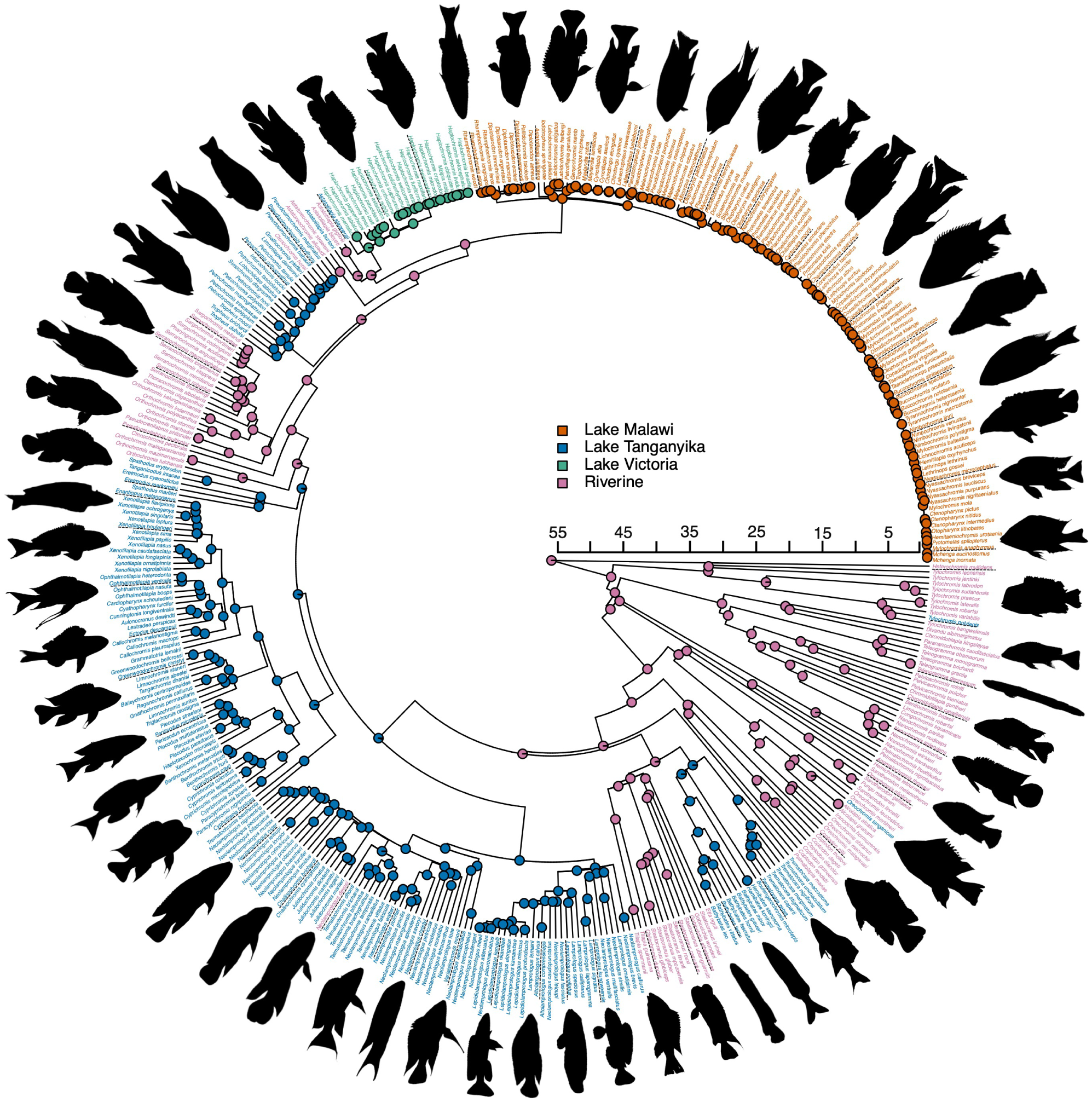
Ancestral character estimation of water system occupancy estimated by maximum likelihood. Uncertainty in node prediction is shown in the form of a pie chart for each node from 1000 simulations (see Methodology). Tips are coloured according to their water system occupancy. Silhouettes of select extant species are shown. Scale is in millions of years. See Supplementary Materials for image credits.

**Supplementary Figure 3.**
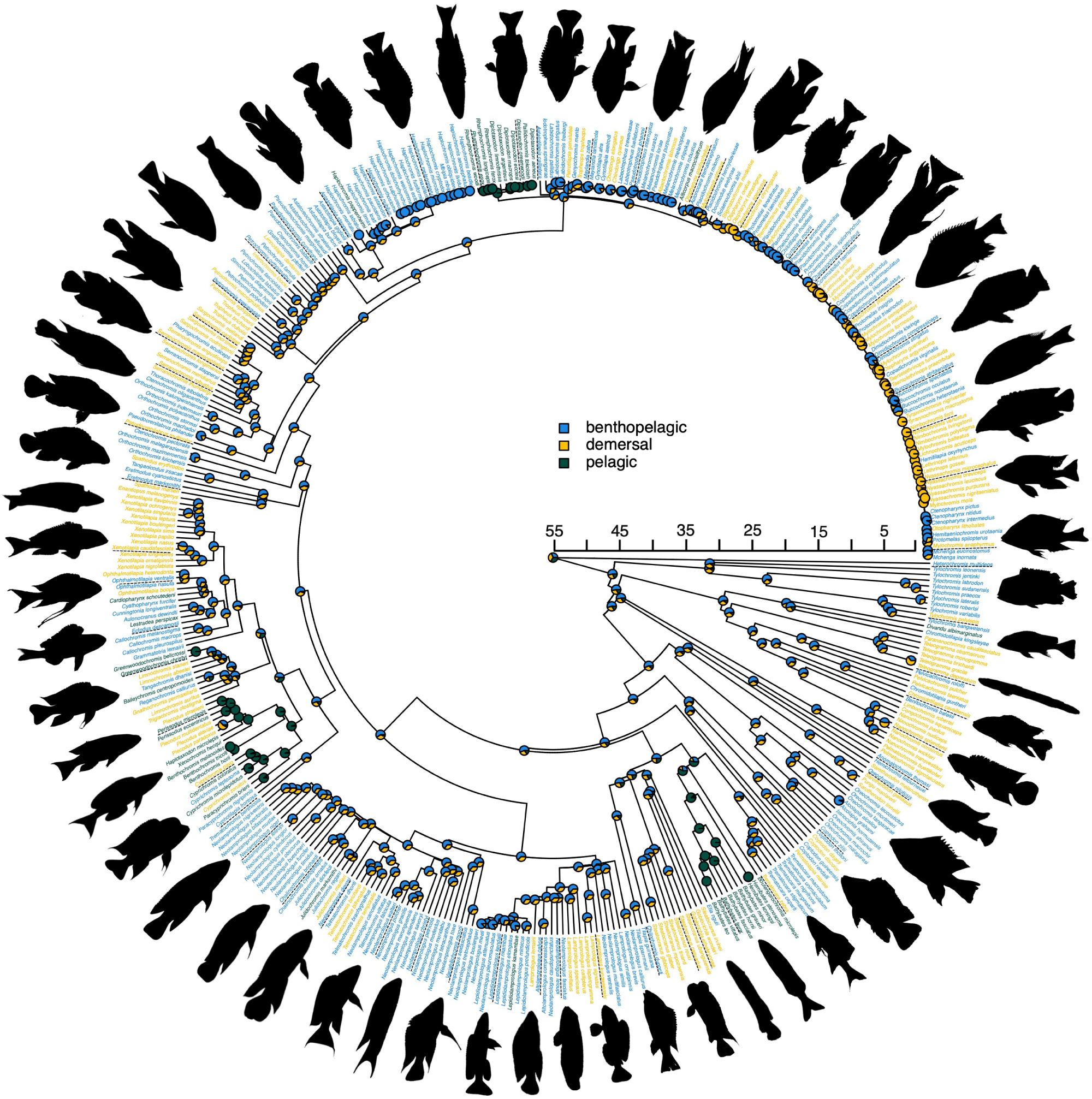
Ancestral character estimation of occupation along benthic-pelagic axis estimated by maximum likelihood. Uncertainty in node prediction is shown in the form of a pie chart for each node from 1000 simulations (see Methodology). Tips are coloured according to presence along the benthic-pelagic axis. Silhouettes of select extant species are shown. Scale is in millions of years. See Supplementary Materials for image credits.

**Supplementary Figure 4.**
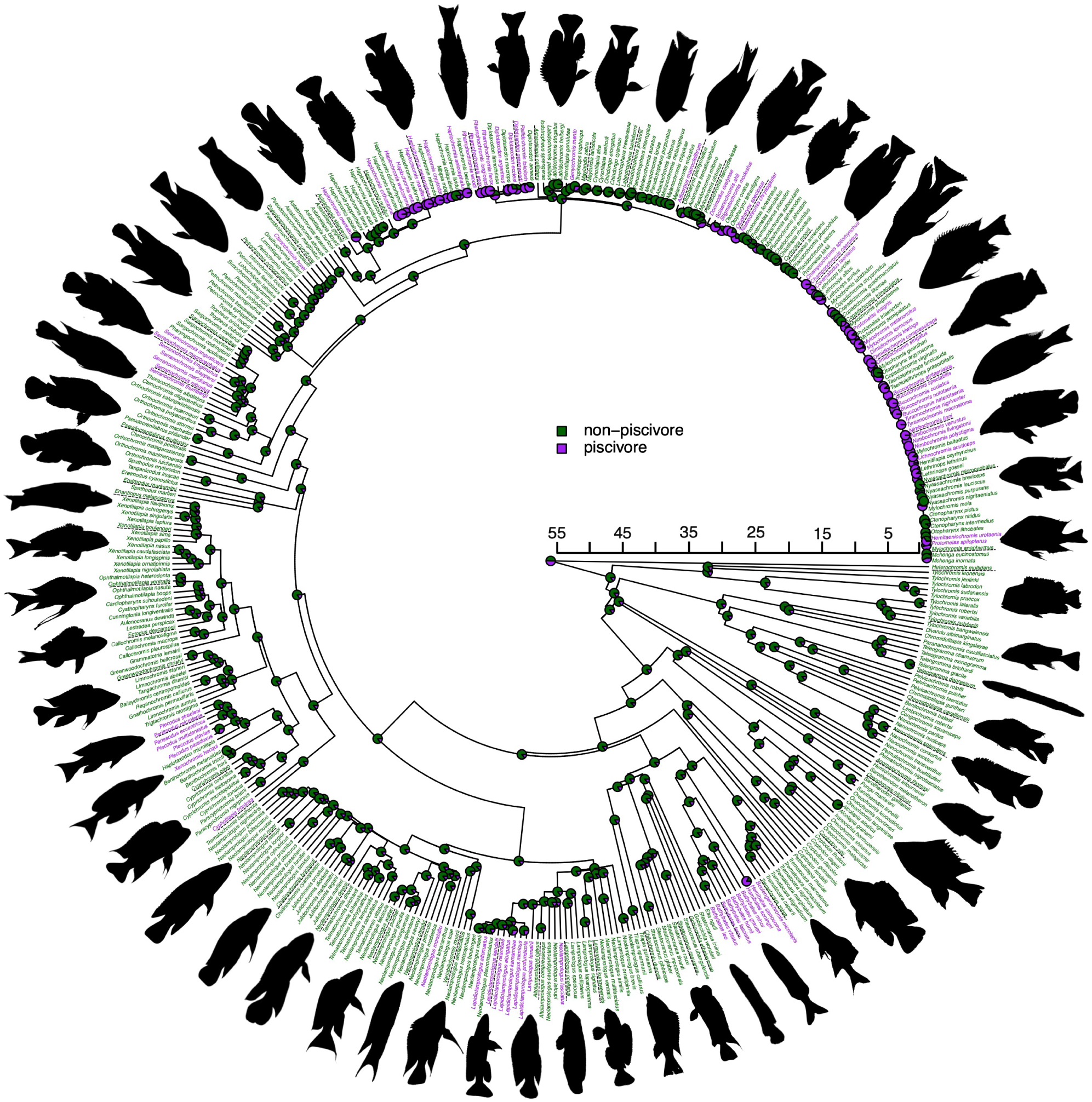
Ancestral character estimation of piscivory estimated by maximum likelihood. Uncertainty in node prediction is shown in the form of a pie chart for each node from 1000 simulations (see Methodology). Tips are coloured according to whether they are piscivorous or not. Silhouettes of select extant species are shown. Scale is in millions of years. See Supplementary Materials for image credits.

**Supplementary Figure 5.**
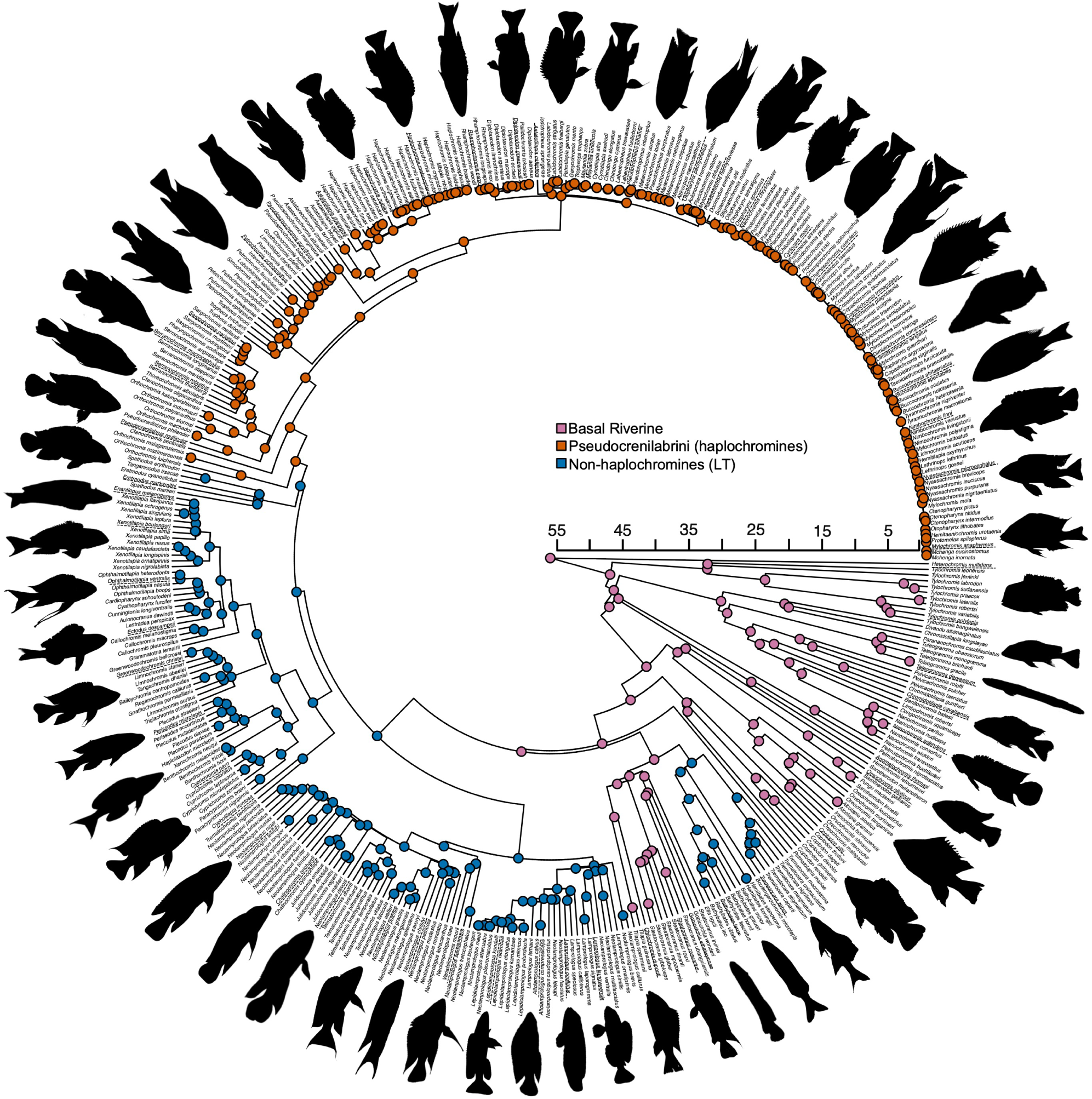
Ancestral character estimation of Pseudocrenilabrini (haplochromines) estimated by maximum likelihood. Uncertainty in node prediction is shown in the form of a pie chart for each node from 1000 simulations (see Methodology). Tips are coloured according to whether they are haplochromines or belong to tribes specific to Lake Tanganyika or are exclusively riverine taxa. Silhouettes of select extant species are shown. Scale is in millions of years. See Supplementary Materials for image credits.

**Supplementary Figure 6.**
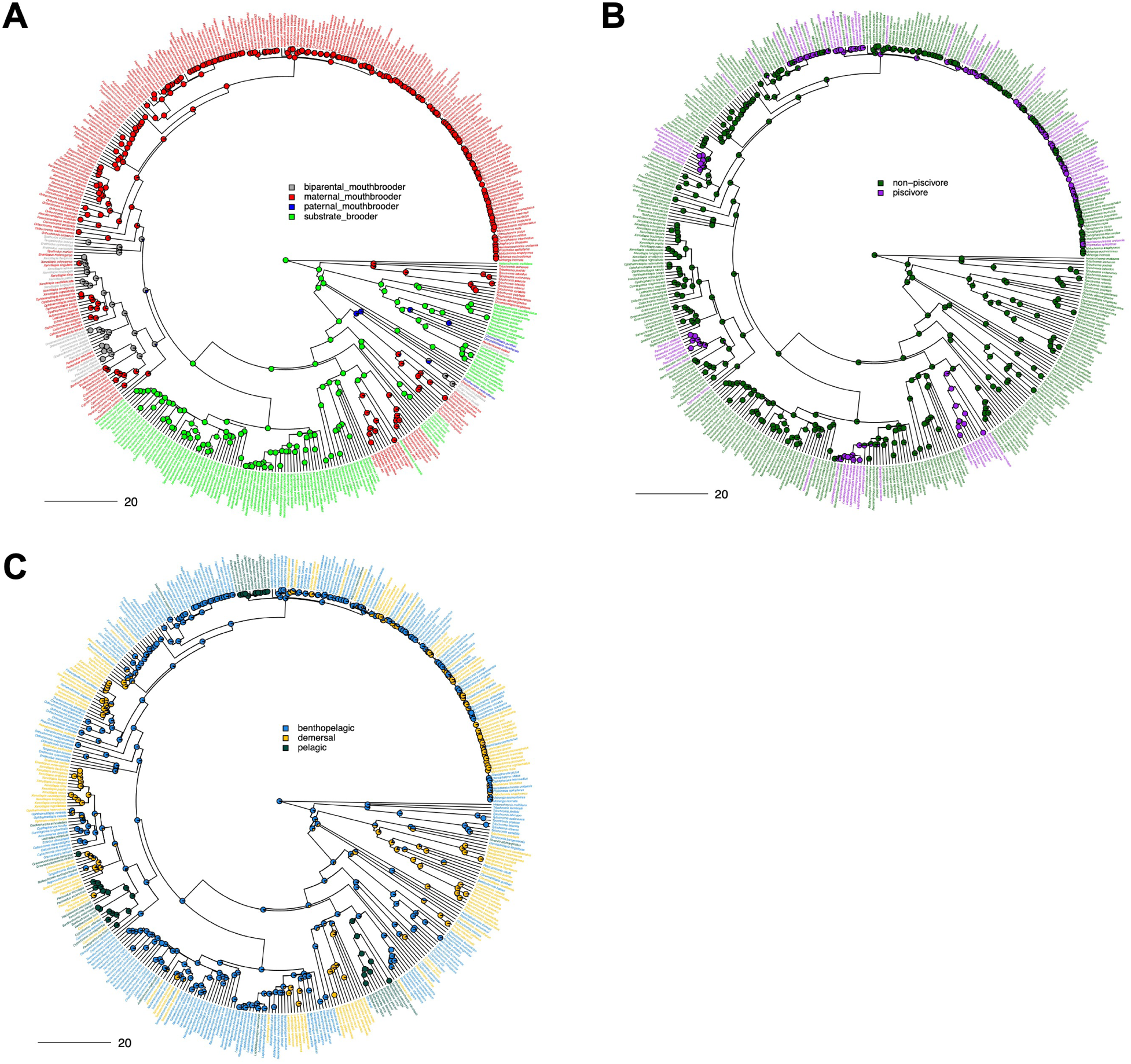
Ancestral character estimation for common ancestor of African cichlids. (A) Brooding behaviour; (B) piscivory; (C) presence along the benthic-pelagic axis. Uncertainty in node prediction is shown in the form of a pie chart for each node from 1000 simulations with branch lengths set to 1 (see Methodology). Branch lengths were replaced to the correct lengths in millions of years for visualisation. Tips are coloured according to the respective ecological group of the extant species. See Supplementary Materials for image credits.

**Supplementary Figure 7.**
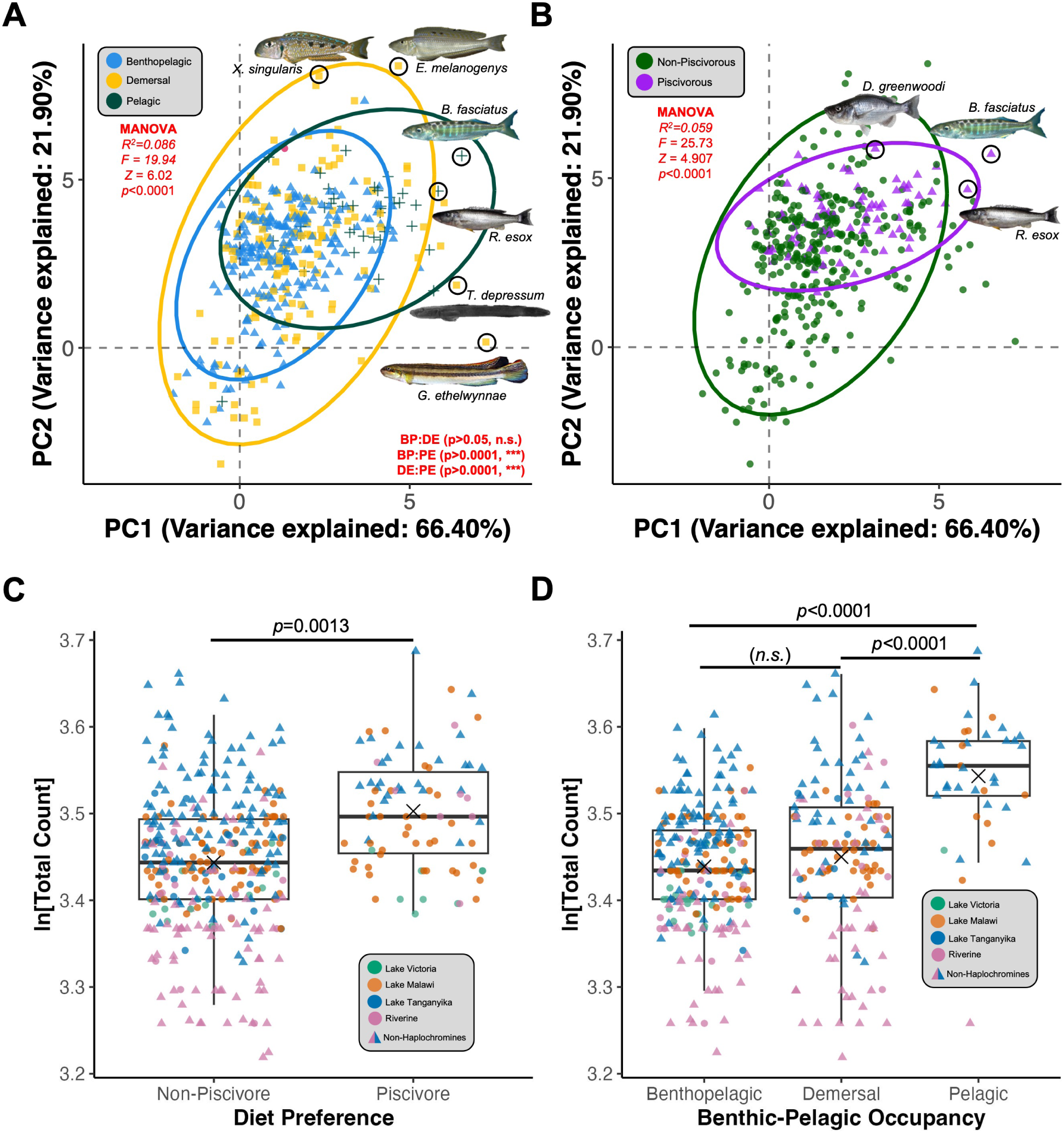
Piscivory and benthic-pelagic axis occupation is correlated with changes in axial morphology. Phylogenetic PCA plotting PC1 against PC2. The variance explained by PC1 and PC2 is indicated on the axes, see Figure 2A for loadings. Species have been grouped according to their occupation along the benthic-pelagic axis (A) or piscivory (B). Spheres around each group cluster represent the 95% CI. Distributions of ln[Total Count] by depth preference (C) and piscivory (D) for all species. The water system each species belongs to is indicated by the key. Means for each group are indicated with a black cross. Means were compared with a phylogenetic ANOVA (see Methodology), the significance of each mean difference is indicated.

**Supplementary Figure 8.**
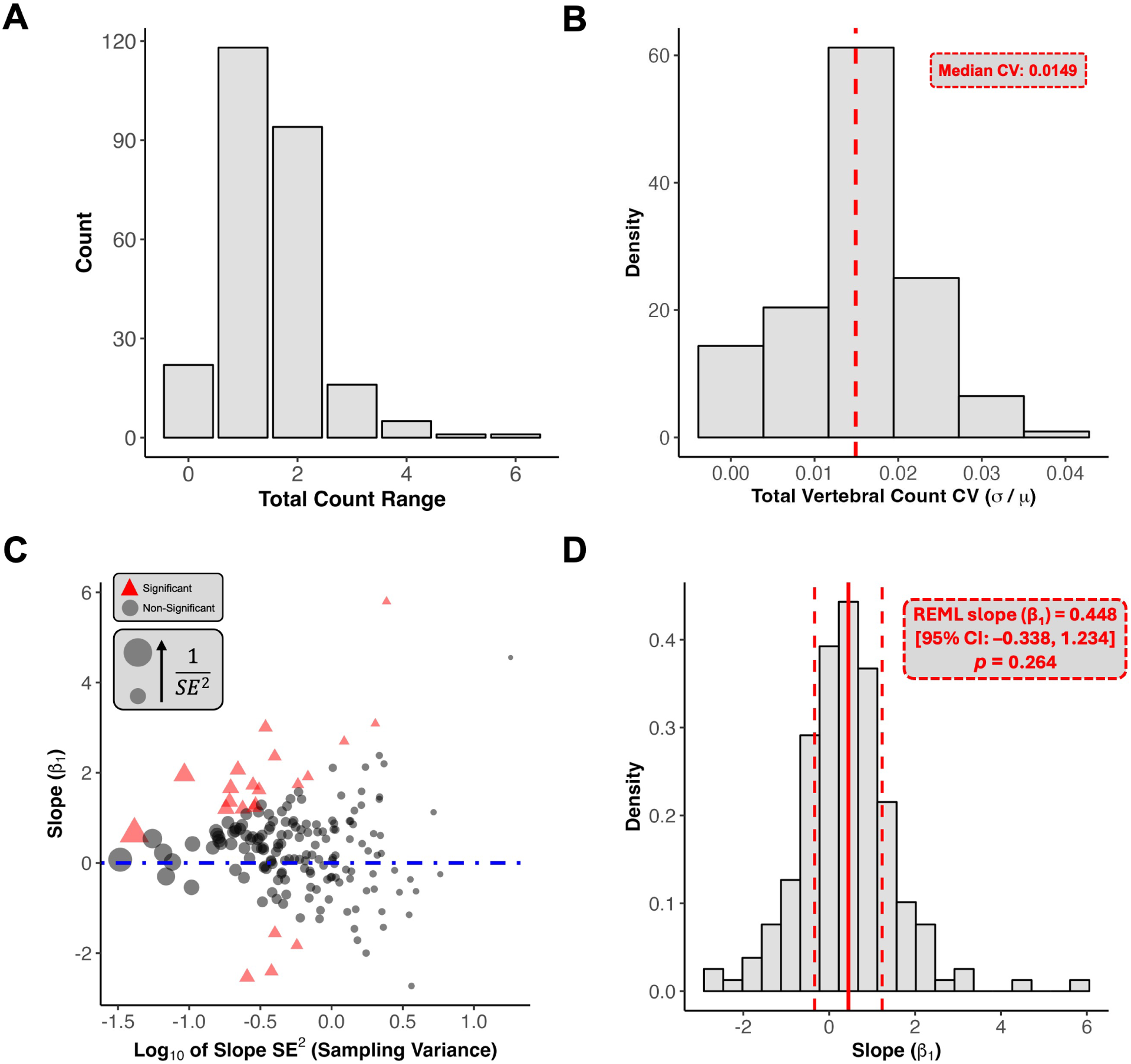
Total vertebral count intraspecific variation is low and does not scale predictably with body elongation. (A) Distributions of total vertebral count range for species with at least 10 individuals (n=257). (B) Distribution of total count coefficient of variation (CV). The CV represents the proportion of the mean total count that tends to vary within a species, the median for 176 species (*n ≥* 10, retained on phylogeny) is indicated. (C) Scatterplot of intraspecific regression slopes (*β*_1_) of ln[Total Vertebral Count] on ln[Length]-ln[Depth] against their respective sampling variances. Point sizes are scaled according to the weights used in the meta-analysis; larger points reflect more precise estimates and thus greater weight. Red triangles indicate slopes significantly different from zero (*p <* 0.05). A dot-dashed blue line indicates a slope of zero. (D) Histogram showing the distribution of intraspecific slopes across all species retained on the phylogeny with at least ten individuals (n=176). The solid red line shows the restricted maximum likelihood (REML) estimate of the mean slope; dashed red lines denote the 95% confidence interval. Note that the confidence interval overlaps zero.

**Supplementary Figure 9.**
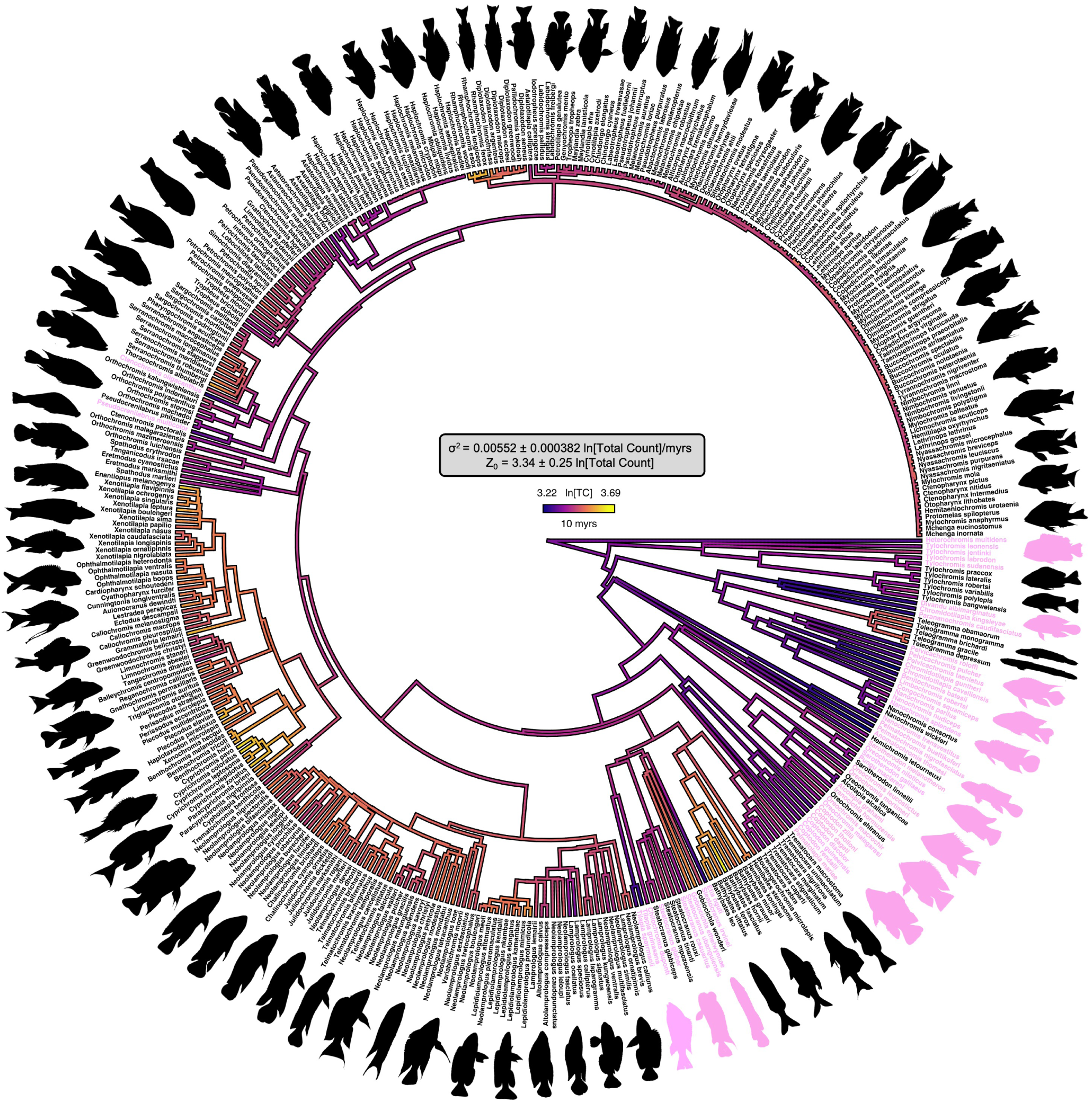
Ancestral trait estimation of ln[Total Count] in Pseudocrenilabrinae. A single rate Brownian motion model was fit to the subfamily tree as per the methodology. *x̅ ±* SD is shown for the estimated stochastic rate (*σ*^2^) and the root trait value (*Z*_0_). Species present within the distinct riverine axial morphospace are shown as pink tips, the body shape of some of these species is also shown as pink silhouettes. Note that the majority of the species within this distinct region of axial morphospace are predicted to have undergone losses in ln[Total Counts] (dark blue) and are relatively basal on the tree.

**Supplementary Figure 10.**
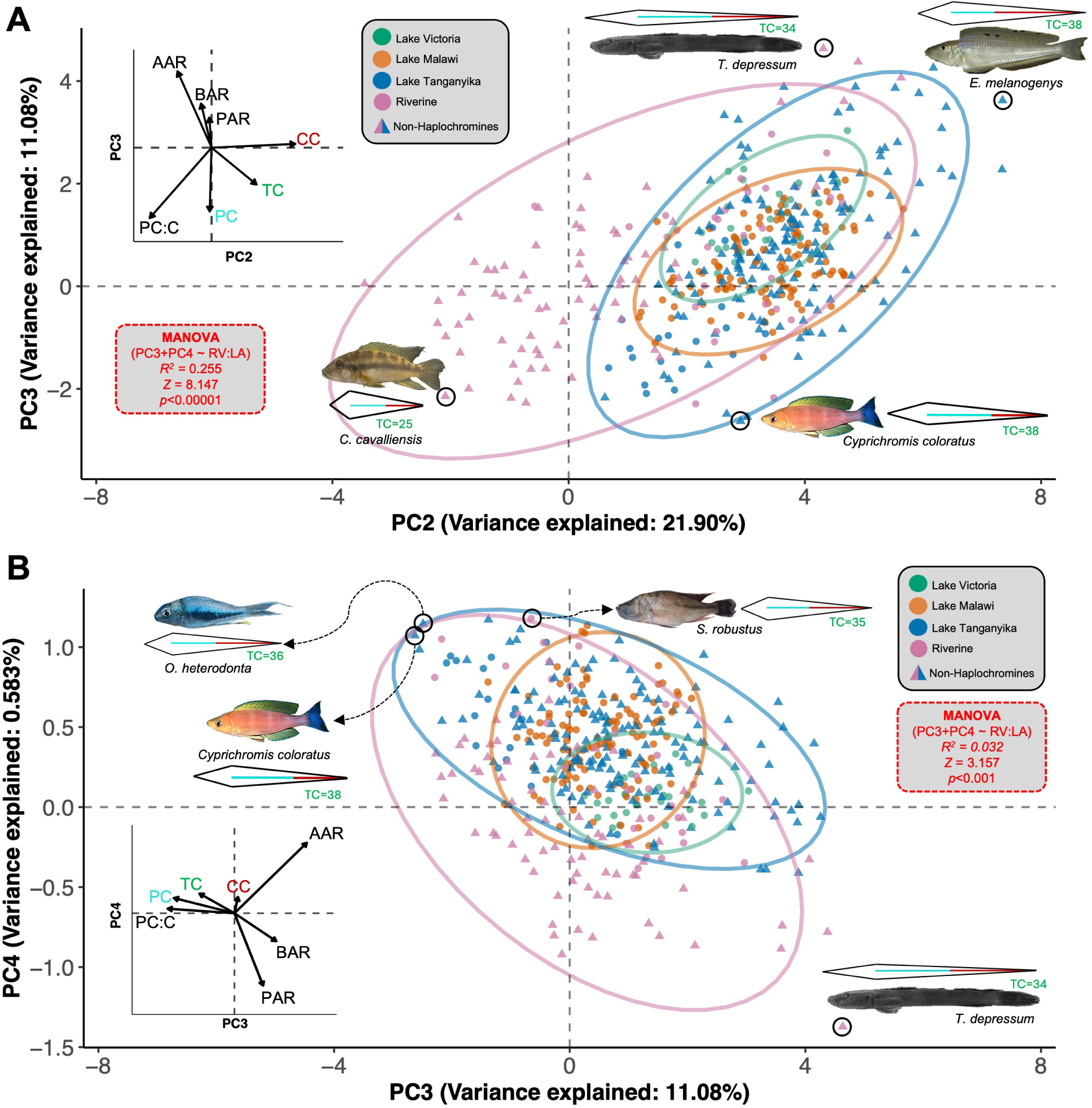
Asymmetrical morphospace occupation between riverine and lacustrine taxa is present at higher PCs. Phylogenetic PCA plotting PC2 against PC3 (A) and PC3 against PC4 (B). The variance explained by each PC is indicated on the axes. A loading plot on the same axes is displayed for both plots. Species representing the extremes of each PC are indicated, along with a graphical representation of their axial phenotype. Modal total vertebral counts for each species are indicated in green. Spheres around each water system cluster represent the 95% CI. MANOVA results are indicated in a dashed red box and test for significant differences in the occupation in PC2+PC3 or PC3+PC4 between riverine (RV) and lacustrine (LA, Lake Victoria, Lake Malawi and Lake Tanganyika) species.

**Supplementary Figure 11.**
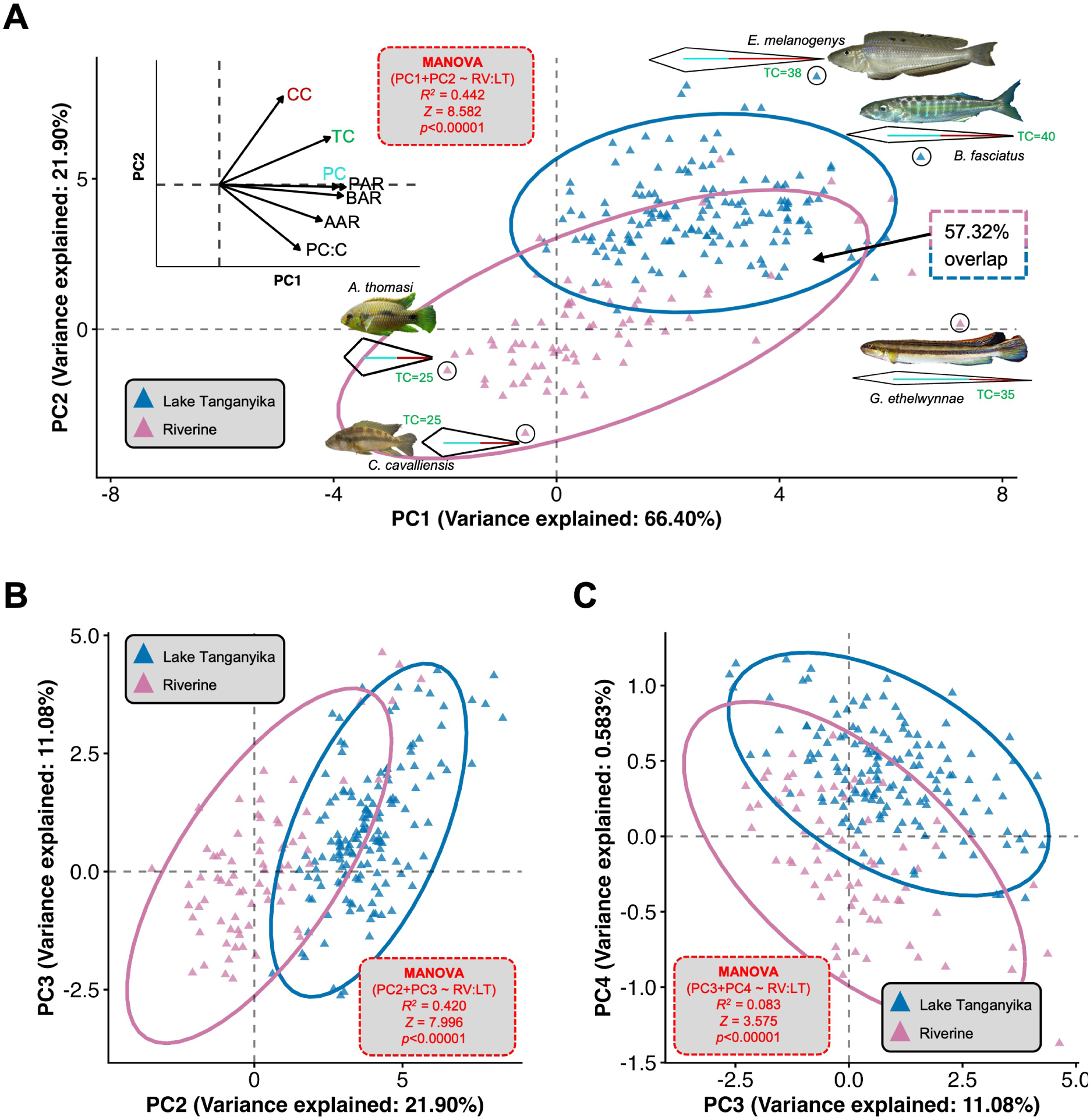
Removal of haplochromines increases distinctness of axial morphospace occupation. Phylogenetic PCA plotting PC1 against PC2 (A), PC2 and and PC3 (B) and PC3 against PC4 (C) in the absence of haplochromines (including Tropheini). The percentage of non-haplochromine Lake Tanganyikan axial morphospace (57.32%) occupied by riverine taxa is indicated for PC1 and PC2 only. The variance explained by each PC is indicated on the axes. A loading plot on the same axes is displayed for (A), see Supplementary Figure 10 for loadings for (B) and (C). Species representing the extremes of PC1 and PC2 are indicated, along with a graphical representation of their axial phenotype. Note that these match with Figure 3A, besides the now removal of Rhamphochromis esox from Lake Malawi. Modal total vertebral counts for each species are indicated in green. Spheres around each water system cluster represent the 95% CI. MANOVA results are indicated in a dashed red box and test for significant differences in the axial morphospace occupation between riverine (RV) and Lake Tanganyika (LT) species.

## AUTHOR CONTRIBUTIONS

C.V.B, B.V and R.B conceived the study. C.V.B. collected the data and performed the data analysis. C.V.B, B.V and R.B helped interpret results. E.D.R helped with digitisation of x-rays. F.R., M.K.O, C.V.B and N.V provided x-ray images of specimens. M.S., F.R. and W.S. and provided access to their respective collections for data collection. M.M. collected the CAMZM specimens in the dataset. C.V.B wrote the manuscript. B.V. and R.B. edited the manuscript. All authors reviewed the manuscript.

## ACKNOWLEDGMENTS

This research was supported by a Biotechnology and Biological Sciences Research Council (BBSRC) studentship (Grant No. 2445747), the John Fell Fund at the University of Oxford (Grant No. 0009780), a European Research Council Starting Grant (No. 101163722, Counts), and grants from the Swiss National Science Foundation (SNSF; Nos. 176039 and 208002). We thank Richard Durbin for the *µ*CT-scans of the specimens in the CAMZM collection used in this study that were *µ*CT-scanned at the Cambridge Biotomography Centre. Specimens were collected ethically under prescribed permits, and the results and data are published under an Access and Benefit Sharing agreement with the Government of Malawi. We acknowledge the contributions of Hannes Svardal, Karl Svardal, Richard Zatha, Bosco Rusuwa, George Turner and colleagues, and the Malawi Department of Fisheries and the Government of Malawi for their assistance in the collection of samples. We thank Ad Konings, George Turner and Adrian Indermaur for the images used in the figures. Thank you to the fishes whose lives were sacrificed for this work.

