## Supplementary Methods for "Replicated radiations reveal temporally structured macroevolutionary dynamics"

### Collating an X-Ray Library

We collated a library of 2D-lateral images of 4544 individuals of Pseudocrenilabrinae. Of the 4544, 3633 were from previously published studies or generated ourselves. This included: 2D lateral X-rays (n=2183) of cichlids from the Lake Tanganyika basin (Ronco et al., 2021); whole-body 2D lateral x-rays of specimens part of the American Museum of Natural History (AMNH) ichthyology collection (n=764); haplochromine riverine specimens part of the research collection at the University of Basel (n=310); 132 individuals of 2D whole-body lateral x-rays generated by Michael Oliver (Oliver, 2024); 131 Lake Victoria region haplochromines generated by Nathan Vranken; lateral images generated from whole-body volume renderings or 3D models of the whole skeleton of $\mu$CT-scans (n=113) of Lake Malawi haplochromines (Bucklow et al., 2024) and whole-body volume renderings from $\mu$CT-scans (n=9) of Lake Malawi cichlids from the Durbin Lab (University of Cambridge) that are part of the Museum of Zoology, University of Cambridge (CAMZM) collection. The remaining images (n=911) were collated from a manual search for each of the 166 currently recognised genera (see below) that constitute Pseudocrenilabrinae in multiple museum collections, including the AMNH ichthyology collection using their own data portal, the Natural History Museum, London (NHMUK) data portal (Scott et al., 2019) and all the museum collections associated with GBIF. Species names were initially taken from the museum record for the specimen(s) and the recorded species name was checked on FishBase (Froese and Pauly, 2000) and updated to the senior synonym, if required. For Lake Malawi haplochromines, this was also supplemented by the curated species list available on malawi.si.

### Vertebral Count and Body Shape Measures

Individual cropped images were manually landmarked using the multi point tool in FIJI (Schindelin et al., 2012), a GUI for ImageJ (Schneider et al., 2012). For landmarks used see Supplementary Figure 1A. Vertebral centra were counted anterior to posterior, starting with the anterior-most, ribless vertebrae, finishing and including the urostyle, the vertebra that articulates with the caudal-fin (Woltering et al., 2018; Di'Biagio et al., 2022). The total number of centra was then recorded as the total vertebral count of that specimen. Where fused centra were visible, we instead counted the number of neural spines (Oliver, 2024) (Supplementary Figure 1B). The precaudal count included the two most anterior vertebrae, which are not associated with pleural ribs, as well as the rib-associated precaudal vertebrae (with zygapophyses). The caudal count included all vertebrae posterior of the last rib-bearing precaudal vertebrae which have haemal spines (and therefore a haemal arch) (Ford,

1937). The presence of '*transitional*' vertebrae (See Supplementary Figure 1C, D), bearing both precaudal and caudal vertebral morphology (De Clercq et al., 2017) made defining the final precaudal and the first caudal vertebrae difficult. '*Transitional*' vertebrae were defined as caudal unless fully formed pleural ribs were present (Supplementary Figure 1C). The presence of a haemal spine, and therefore haemal arch, overrode the classification regardless of pleural rib presence (Supplementary Figure 1D). We also measured three linear measures to quantify body shape changes: a whole body aspect ratio quantifying the relationship between anteroposterior length and dorsoventral depth of the whole body, an anterior body aspect ratio (broadly quantifying elongation of the head) and a posterior body aspect ratio (quantifying postcranial elongation), using previously described landmarks (Ronco et al., 2021), see Supplementary Figure 1A. Since many of the radiographs lacked scale bars, we calculated raw pixel lengths and widths from the coordinates of the landmarks for the body aspect ratio and calculated a ratio between the log-transformed measures. 71 specimens were filtered from the dataset because of missing craniums, pectoral fin rays etc.

### Taxonomic Considerations and Defining Water Systems

We wanted to sample the Pseudocrenilabrinae as widely as possible. Therefore, we focused on maximising the sampling of every tribe and genus within these tribes. We used the detailed discussion of Pseudocrenilabrinae taxonomy of Astudillo-Clavijo et al. (2023) to collate a list of genera that belong to each tribe not endemic to Lake Tanganyika including: Chromidotilapiini, Coptodonini, Gobiocichlini, Hemichromini, Heterochromini, Oreochromini, Pelmatochromini, Pelmatolapiini, Steatocranini, Tilapiini and Tylochromini. For Lake Tanganyika endemic tribes, we instead used the detailed taxonomic discussion of endemics and native species to Lake Tanganyika (Ronco et al., 2020), as well as the phylogeny constructed by Ronco et al. (2021), which resolved the phylogenetic relationships between all tribes of Pseudocrenilabrinae that are endemic or native to Lake Tanganyika or the wider basin, respectively. Tribes endemic or native to Lake Tanganyika included: Bathybatini, Benthochromini, Boulengerochromini, Cyphotilapiini, Cyprichromini, Ectodini, Eretmodini, Lamprologini, Limnochromini, Perissodini, Trematocarini, Tropheina (Lake Tanganyika endemic haplochromines) and haplochromines endemic to the wider basin, but not part of the Lake Victoria or Malawi radiations. All species belonging to any of the Lake Tanganyika endemic tribes were classified as belonging to the Lake Tanganyika water system, besides *Telmatochromis devosi*, formerly *Neolamprologus* (Indermaur et al., 2024), a Lamprologini, a species found in the lower Malagarasi river, an inlet of Lake Tanganyika (Schelly et al., 2003). We also note that Lamprologus now refers exclusively to lamprologines occurring outside Lake Tanganyika following recent taxonomic reassignment (Alda et al., 2025); however, we retain the historical nomenclature here for consistency. In addition, we did not sample any of these (now) riverine lamprologines and thus they were not included in any of the analyses.

All haplochromine genera belonging to the Lake Malawi radiation, were classified as such, including species endemic to satellite lakes of Lake Malawi (Turner et al., 2019). *Astatotilapia calliptera* is distributed across the rivers flowing eastward to the Indian Ocean, from the Rovuma River in the north, to the Save River in the south (Turner et al., 2021) and despite its exceptionally wide distribution and riverine ecology, it clusters phylogenetically within the Lake Malawi radiation (Malinsky et al., 2018). Therefore, all specimens of *Astatotilapia calliptera* were categorised as being part of the Lake Malawi system and not riverine. Similar exceptions were made for *Astatotilapia burtoni* and *Astatotilapia stappersii*, non-Tropheini haplochromines native to Lake Tanganyika and its wider catchment, including its connecting waterways (Turner et al., 2021). Other *Astatotilapia* species included: *A. gigliolii*, part of the '[Great] Ruaha [River] catchment' that belongs to a sister lineage to the haplochromine radiations of Lake Malawi and Lake Victoria (Svardal et al., 2020); *A. bloyeti*, a species distributed in the Wami River system, the Malagarasi, the Pangani and the catchments of Lake Manyara and Eyasi and associated lakes; *A. paludinosa*, whose distribution overlaps with that of *A. bloyeti* (Turner et al., 2021) were all classified as 'Riverine' species. Haplochromines that do not belong to either the Lake Malawi or Lake Victoria radiations were classified as riverine. For example, *Serranochromis robustus* belongs to one four genera referred to as 'serranochromines' (Greenwood, 1993). *Serranochromis robustus* is native but not endemic to Lake Malawi (Turner et al., 2019). Phylogenetic analyses show that serrachromines do not cluster within the Lake Malawi radiation (McGee et al., 2020; Astudillo-Clavijo et al., 2023), and members of the assemblage are found throughout East African river systems (Thorstad et al., 2005). This, along with evidence suggesting that the group originated in a now extinct lake before colonising nearby rivers (Joyce et al., 2005), supports their classification as riverine.

All haplochromine species belonging to the Lake Victoria Region Superflock (LVRS) were grouped into the 'Lake Victoria' clade. This includes all haplochromine species endemic to Lakes Victoria, Albert, Edward, George and Kivu (Meier et al., 2017), as well as Lake Kyoga whose endemics were not considered by Meier et al. (2017). However, Lake Kyoga is nonetheless directly connected to Lake Victoria by the White (Victoria) Nile (Mwanja et al., 2001) and more recent evidence suggests that the Lake Kyoga endemics are indeed part of the LVRS (Meier et al., 2023). Only eight total specimens (three species) were from Lake Kyoga, including the holotype of *Haplochromis worthingtoni*, NHMUK 1929.1.24.334; six syntypes of *Haplochromis latifasciatus* from NHMUK 1929.1.24.335-339; and a specimen of *Paralabidochromis sp.* "black" from the Museum of Comparative Zoology (MCZ), MCZ:Ich:137961, the latter of which was not present on the phylogeny and not included in any phylogenetic analysis.
